# Clinical isolates of *Fusobacterium nucleatum* display strain-specific virulence and modulation by indole derivatives

**DOI:** 10.1101/2025.05.16.654558

**Authors:** Colin Scano, Ankan Choudhury, Macarena G. Rojo, Jessalyn Hawkins, Lauren Crowhurst, Greg Zaharas, Ramon Lavado, Leigh Greathouse

**Affiliations:** Biology Department, Baylor University; Robbins College of Health and Human Sciences, Baylor University; Environmental Science, Baylor University

**Keywords:** *Fusobacterium nucleatum*, biofilm, indole, tryptophan metabolism, indole derivatives

## Abstract

Pathogenic bacteria adapt to distinct disease environments, but whether these adaptations create therapeutic vulnerabilities remains unclear. *Fusobacterium nucleatum* has emerged as a key microbial player in colorectal cancer (CRC), yet its strain-specific virulence mechanisms remain poorly defined. In this pilot study of 16 clinical *F. nucleatum* isolates from CRC patients (n=6), Crohn’s disease patients (n=6), colon of healthy individuals (n=3), and an oral lesion (n=1), a subset of CRC-derived strains produced 3–4-fold higher levels of endogenous indole. Exogenous indole treatment differentially affected growth and biofilm formation, with some strains increasing biofilm despite growth inhibition. Notably, sensitivity to exogenous indole was independent of endogenous production and revealed that isolate 7-1 (EAVG002), a member of the tumorigenic *Fna* C2 clade, was uniquely hypersensitive to I3CA- and IPA-mediated stress. Invasion assays further showed that indole and its derivatives (I3A, I3CA) reduced invasion of a highly indole-tolerant CRC-derived isolate (SB-CTX3Tcol3) into CRC cells by ∼50%, comparable to antibiotic treatment. Furthermore, in two CRC cell lines, exposure to indole or indole derivatives resulted in substantial variability in adherens junction and tight junction transcript levels, with I3A having the strongest effect on tight junction (*CLDN1, CLDN7*) transcripts. Collectively, these findings reveal profound strain-level heterogeneity and indole derivative effects, highlighting vulnerabilities that could enable precision therapeutic targeting of pathogenic *F. nucleatum* populations within the host environment while preserving beneficial commensals.

## INTRODUCTION

The discovery that *Fusobacterium nucleatum (F. nucleatum)* selectively colonizes colorectal cancer (CRC) (1,2) transformed our understanding of host-microbe interactions in cancer, revealing that pathogenic bacteria not only respond to disease environments but may actively drive oncogenesis. While these *F. nucleatum* populations are thought to originate from the oral cavity, it remains unclear whether disseminated strains retain their original virulence programs or undergo niche-specific adaptation to the tumor environment (3). Recent pangenomic analysis suggests that virulence determinants and functional clusters in *F. nucleatum* are largely strain-specific rather than subspecies-specific, highlighting the genomic plasticity of individual isolates to adapt to distinct niches and optimize fitness (4).

*F. nucleatum* is selectively enriched in invasive bacterial biofilms associated with CRC (5), a pathogenic mechanism that may also be mirrored in inflammatory bowel and Crohn’s disease (6,7). These invasive bacterial biofilms are carcinogenic in mouse models and correlate with reduced E-cadherin expression and increased crypt cell proliferation, directly linking bacterial biofilm to disease progression (5,8,9). Though *F. nucleatum*’s role in Crohn’s disease is still under investigation, CD-derived isolates—specifically the tumorigenic *Fna* C2 strain 7-1 (EAVG002)—exhibit invasive and pro-inflammatory phenotypes that mirror those of CRC-derived strains (7,10,11).

Beyond basic biofilm formation, *F. nucleatum* deploys an arsenal of outer membrane proteins and adhesins (including RadD, Aid1, and FadA), alongside the major structural porin FomA, that mediate both polymicrobial interactions and host cell binding (12–17). Furthermore, adherence to host cells enables *F. nucleatum* to trigger pro-inflammatory signaling pathways. The best-characterized adhesin, FadA, binds to E-cadherin to facilitate invasion (18), activate β-catenin signaling (15,19), and exacerbate DSS-induced colitis (20). Similarly, FomA serves as a critical anchor for epithelial adherence and immune modulation (16,21). Yet a critical knowledge gap remains: what environmental signals regulate these diverse virulence programs in the diseased environment?

Identifying mechanisms regulating niche adaptation, including biofilm formation, virulence factor expression, and host cell invasion, is central to understanding *F. nucleatum’s* pathogenicity (5,15,22). One candidate regulator of niche adaptation is indole, a chemical effector in enteric communities that modulates biofilm formation and virulence across various pathogens (23). While early studies indicated that indole enhances biofilm formation in the oral-derived *F. nucleatum* reference strain ATCC 25586, it remains unknown whether this response is conserved across clinical isolates (24). Recent meta-analyses of fecal microbiome data indicate that tryptophan biosynthesis and metabolism are significantly enriched in CRC tumor samples compared to healthy colon controls **(Supplementary Figure 1)** (25–27) and increased indole levels can contribute to increased gut inflammation (28). Beyond these broad shifts, recent evidence has confirmed that specific indole derivatives produced by *F. nucleatum* can drive pro-carcinogenic pathways (29,30). However, it remains unclear how the production of indole and its derivatives regulate *F. nucleatum* virulence, and whether these metabolites act as niche-specific signals.

In this pilot investigation, we hypothesized that *F. nucleatum* isolates from distinct disease environments harbor unique adaptations manifesting as differential responses to indole. We specifically examined whether a strain’s phenotypic response to exogenous indole is independent of its endogenous production profiles, and how specific indole derivatives affected pathogenic behavior. Overall, our study identified strain-specific responses to indole and indole derivatives and identified metabolic vulnerabilities that may be exploited for selective therapeutic suppression.

## MATERIALS AND METHODS

### Bacterial Anaerobic Culture Conditions

*F. nucleatum* isolates were cultured from glycerol stocks on anaerobically reduced Columbia blood agar (5% sheep blood) or in Columbia broth supplemented with 0.5 g/L L-cysteine, 5 mg/L hemin, and 10 µg/L vitamin K (CB). Cultures were incubated at 37°C for 48–72 hours in a Coy Laboratories anaerobic chamber (90% N_2_, 5% CO_2_, 5% H_2_).

### Quantification of total indole production with Kovacs reagent

Total indole production was quantified using Kovac’s reagent. Stationary phase cultures were adjusted to 0.3 OD_600_ and inoculated into four media types: CB, Brain Heart Infusion (BHI), or CB supplemented with 5 mM tryptophan or 1% glucose. Supernatants collected at 12-hour intervals (up to 72 h) were reacted with an equal volume of Kovac’s reagent for 5 minutes. Absorbance (OD_560_) was measured using a Sunrise Tecan reader, and concentrations were determined via a 0–2000 µM standard curve. Data represent three biological replicates per isolate across each media condition.

### Quantification of indole derivatives by HPLC

Starter cultures were adjusted to 0.3 OD_600_ and inoculated into CB for 48 h. Supernatants were collected via centrifugation and analyzed using an UltiMate 3000 HPLC system with an RS fluorescence detector (Thermo Fisher Scientific) according to previously described methods (31,32). Separation was performed on an Acclaim™ VANQUISH C-18 column (2.2 μm, 150 × 2.1 mm) at 30°C with a flow rate of 0.5 mL/min. The mobile phase utilized a linear gradient of eluent A (50 mM KH_2_PO_4_, pH 3.3) and eluent B (70:30 eluent A:acetonitrile): 20% B (0–2 min), 20% to 75% B (2–10 min), 100% B (10–15 min), and 20% B (15–20 min). Native fluorescence was monitored at excitation/emission wavelengths of 270/350 nm. Targeted compounds (indole (CAS# 120-72-9), IPA (CAS# 830-96-6), IAA (CAS# 87-51-4), I3A (CAS# 487-89-8), and I3CA (CAS# 771-50-6)) were identified via authentic standards, confirmed by LC-MS, and quantified by peak area integration. Method linearity and system suitability (retention time stability, resolution) were monitored throughout.

### Quantification of Biofilm Biomass

Biofilm biomass was quantified in 96-well plates via crystal violet (CV) staining (24). Briefly, biofilms were washed, air-dried, and fixed with methanol (37°C, overnight). Fixed biofilms were stained with 0.1% CV for 1 minute, washed, and destained with 30% acetic acid. Absorbance (A_560_) was measured using a Sunrise Tecan microplate reader following a 10-second orbital shake.

### Quantification of F. nucleatum biofilm over 72 hours

For time-course analysis, isolate cultures were grown to stationary phase, adjusted to 0.3 OD_600_ in CB, and used to inoculate six 96-well plates simultaneously. Two uninoculated wells per plate served as contamination and background controls. Plates were harvested at 12-hour intervals over 72 h; bacterial density was measured (OD_600_) using a Cerillo Stratus reader prior to biofilm quantification as described previously. Six biological replicates were performed per isolate. Biofilm was normalized to the maximum value per timepoint.

### Quantification of Indole-Mediated Growth and Biofilm Alterations

Growth kinetics of 16 isolates were monitored under anaerobic conditions for 72 h (OD_600_ every hour) using a Cerillo Stratus reader. Isolates were treated with five indole derivatives (0.25–2 mM) in triplicate; controls included 0.1% v/v DMSO (vehicle), tryptophan, and chloramphenicol. Following the 72-h kinetic run, the same experimental plates were immediately processed for biofilm quantification as previously described. Statistical significance was determined by one-way ANOVA followed by Tukey’s HSD post-hoc test.

### Statistical Modeling of Indole Sensitivity and Correlation Analysis

Growth (AUC) and biofilm mass were normalized relative to untreated vehicle controls (baseline = 1.0) for each isolate–indole combination **(Supplementary Figure 5, 6, and 7)**. For each isolate–indole combination, signed deviations from the baseline were calculated at each tested concentration (0.25, 0.5, 1.0, and 2.0 mM). To determine overall sensitivity, only the Maximum Signed Deviation (Δ) was retained for each pair; this was defined as the single largest absolute deviation across all concentrations, preserving its original positive or negative sign. Isolates were hierarchically clustered based on Max Δ score profiles across treatments using Euclidean distance and Ward.D2 linkage **(Supplementary Figure 8)**. Statistical significance of the Maximum Signed Deviation (Δ) relative to untreated control was assessed using one-way ANOVA with Benjamini–Hochberg false discovery rate (FDR) correction (p < 0.05, p < 0.01, p < 0.001). To quantify phenotypic response magnitude, growth and biofilm Δ scores were projected against each other and calculated as the Euclidean distance from the untreated origin (0,0): *Responce Magnitude* = √Δ*Growth*^2^ + Δ*Biofilm*^2^. Hyper-responsive isolates were defined as those falling outside a 75% confidence ellipse. The relationship between endogenous indole production and exogenous indole sensitivity was evaluated using Spearman’s rank correlation (ρ) and linear regression.

### Bacterial RNA Isolation and virulence gene qPCR

*F. nucleatum* isolates (initial OD_600_ = 0.3) were treated in triplicate with 1 mM indole derivatives or 0.1% DMSO. After 72 h anaerobic incubation, cells were harvested (8,000×g, 5 min) and lysed using lysozyme (0.5 mg/mL), 0.1 M sodium acetate, and 1% SDS. RNA was isolated via acid-phenol:chloroform extraction at 65°C, purified using Phasemaker tubes, and treated with TURBO DNase. RNA integrity was confirmed by Nanodrop and Qubit assays. cDNA was synthesized from 500 ng of total RNA; no-RT controls confirmed the absence of gDNA. Expression of virulence and metabolic genes (tnaA, aid1, fomA, fadA, radD, **(Supplementary Table 1)**) was quantified using SYBR Green chemistry on a QuantStudio 6 Flex system. Target gene expression was normalized to 16S rRNA and calculated using the 2^−ΔΔC_T_^ method relative to the vehicle control. Primer specificity was confirmed on genomic DNA.

### Phylogenetic Tree Construction

To evaluate the relationship between phylogeny and indole-mediated transcriptional responses, a Maximum Likelihood (ML) tree was constructed using 16S rRNA sequences from 16 isolates **(Supplementary Table 2)**. Sequences were obtained from suppliers, previously published genomic data (33) or generated via Sanger sequencing of an ∼800 bp PCR amplicon (primers 8F/907R) following previously established protocols (34). Sequences were aligned using DECIPHER in R. An initial Neighbor-Joining tree was generated using *phangorn* and optimized via ML for topology, base frequencies, substitution rates, and gamma-distributed rate variation. The tree was rooted with *F. nucleatum* ATCC 25586 and converted to an ultrametric format for direct heatmap alignment using *ggtree* and *aplot* **(Figure 6A)**.

### Predictive Modeling and Statistical Validation

Naive Bayes classifiers were developed to predict clinical origin (CRC, Crohn’s, or Healthy colon) based on indole-induced transcriptional signatures. We compared a quantitative “Fold Change” model (2^−ΔΔC_T_^) against a categorical “Expression Direction” model (thresholds: (Up: >1.25; Down: <0.75; No Change: 1.25–0.75)). Models were trained on a 70% and validated on a 30% test set. Data split was validated using 10-fold cross-validation **(Supplementary Figure 10A).** Performance was evaluated via Accuracy, AUC-ROC, and macro-averaged Precision-Recall curves **(Figure 6E)**. Model calibration was verified through posterior class probabilities **(Supplementary Figure 9C)**, multiclass Brier scores, and faceted calibration curves **(Supplementary Figure 10B-C)**. To ensure feature independence, we calculated Mutual Information **(threshold <0.5; Supplementary Figure 9A)** and Cramér’s V statistics **(Supplementary Figure 9D)**. Feature importance was determined by calculating the Area Under the Curve (AUC) for each gene-class pair to quantify discriminatory power **(Figure 6D).**

### Assessment of Mammalian Epithelial Integrity

Caco-2 and HT-29 cells (gift from Dr. Christie Sayes, Baylor University) were maintained in DMEM supplemented with 10% FBS and 200 μg/mL penicillin-streptomycin at 37°C (5% CO_2_). Cells were seeded in 12-well plates (0.1 x 10^6^/well) and grown to 95% confluence before treatment with 1 mM indole derivatives, 0.1% DMSO (vehicle), or 10 µg/mL LPS (positive control for barrier disruption) for 24 h. Total RNA was extracted and reverse-transcribed, with no-RT controls confirming absence of DNA. qPCR was performed using PowerUp SYBR Green on a QuantStudio 6 system with 250 nM primers **(Supplementary Table 1)**. Relative expression of junctional genes was quantified via the 2*^-ΔΔCt^* method, normalized to β-actin and relative to the vehicle control (DMSO).

### Bacterial Invasion Assay

Caco-2 cells (∼85% confluence) in 24-well plates were incubated in antibiotic- and serum-free DMEM containing 1 mM indole derivatives. Cells were infected with log-phase *F. nucleatum* (isolate SB-CTX3Tcol3) at an MOI of 50:1 for 2 or 4 h. Following infection, monolayers were washed three times with PBS and treated with 200 µg/mL penicillin-streptomycin for 1 h to eliminate extracellular bacteria. After additional washing, cells were lysed with sterile deionized water. Lysates were serially diluted and plated in triplicate on complete medium agar. CFU/mL was calculated after 48 h of anaerobic incubation at 37°C.

### Statistical Methods

All statistical analyses were conducted in R (v4.5.1) using the *tidyverse* package for data curation and *ggplot2* for visualization. Differences across multiple experimental groups were assessed using one-way Analysis of Variance (ANOVA). To control for family-wise error rates during multiple comparisons, ANOVA was followed by Tukey’s Honest Significant Difference (HSD) post-hoc test. Statistical significance was defined a priori at **α** = 0.05. Unless otherwise noted, quantitative data are presented as mean ± standard error of the mean (SEM).

## RESULTS

### Isolates exhibit strain-specific indole production profiles

Using Kovac’s reagent to capture a broad profile of indole-containing metabolites (35), we quantified indole production across 16 clinical *F. nucleatum* isolates over a 72-hour period **(Figure 1A)**. Indole production generally increased after 36 hours, corresponding with the transition from exponential to stationary growth phase. While production was observed for all isolates, there was significant heterogeneity within the isolate groups. Notably, a subset of CRC-derived isolates (SB-CTX3Tcol3, SB-CTX47T, CC7_3JVN3C1, and CC2_3FMU1) exhibited a 3- to 4-fold increase in indole production relative to the other isolates **(Figure 1B)**. This heightened production was consistent across different growth media (**Supplementary Figure 2**). These distinctions were less pronounced when grouped by subspecies, highlighting strain-to-strain differences over classical subspecies classification **(Figure 1C)**. These results demonstrate that, within this cohort of isolates tested, indole production may be strain-specific and partially influenced by clinical origin, independent of subspecies classification.

**Figure 1:**
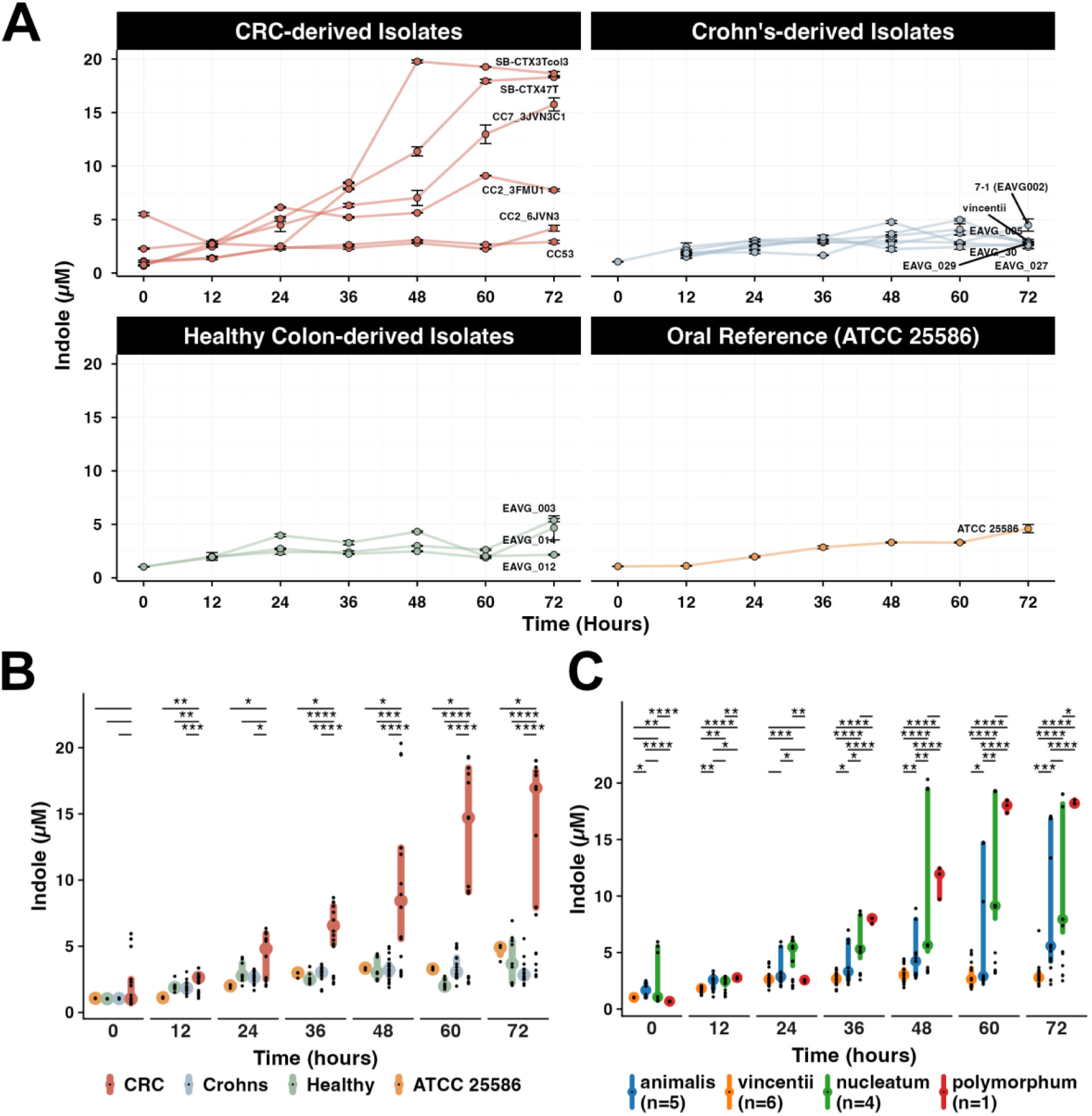
Isolates exhibit strain-specific indole production profiles. Kinetic profile of indole concentration quantified via Kovac’s reagent assay. Values are mean ± SEM per isolate. **(B, C)** Differences in indole levels when isolates are grouped by disease association (B) and subspecies (C). Bars indicate median with IQR. Statistical analysis performed using adjusted ANOVA and Tukey’s HSD comparisons (****p < 0.0001; ***p < 0.001; **p < 0.01; *p < 0.05). Group sizes: CRC (n=6), Crohn’s (n=6), healthy colon (n=3), oral reference strain (ATCC 25586) (n=1). Subspecies groups: *animalis* (n=5), *vincentii* (n=6), *nucleatum* (n=4), *polymorphum* (n=1).

### Isolates exhibit strain-specific indole derivative profiles and metabolic preferences

To characterize the specific metabolic outputs of indole derivatives by individual isolates of *F. nucleatum*, we employed targeted HPLC analysis of five key derivatives that were differentially abundant in CRC patient stool (36). While all targeted derivatives—indole, indole-3-aldehyde (I3A), indole-3-carboxylic acid (I3CA), indole-3-acetic acid (IAA), and indole-3-propionic acid (IPA)—were detected across the cohort, IPA levels remained below the limit of detection in some samples **(Figure 2A**; **Supplementary Figure 3A)**. Notably, a subset of CRC- and Crohn’s-derived isolates exhibited significantly higher production of indole, I3CA, and IAA compared to healthy colon-derived isolates (p < 0.05). Consistent with our earlier findings, subspecies classification did not account for these differences **(Supplementary Figure 3B)**. Correlation analysis and derivative-to-parent indole ratios revealed that isolates possess distinct metabolic signatures or preferences **(Figure 2B)**. Specifically, three CRC-derived isolates (SB-CTX3Tcol3, SB-CTX47T, and CC2_6JVN3) demonstrated a marked preference for I3A production, with I3A/indole ratios (0.52–0.89) substantially exceeding those of other clinical or reference strains **(Figure 2C)**. These data suggest that individual isolates derived from disease environments may harbor specific adaptations in indole derivative production.

**Figure 2:**
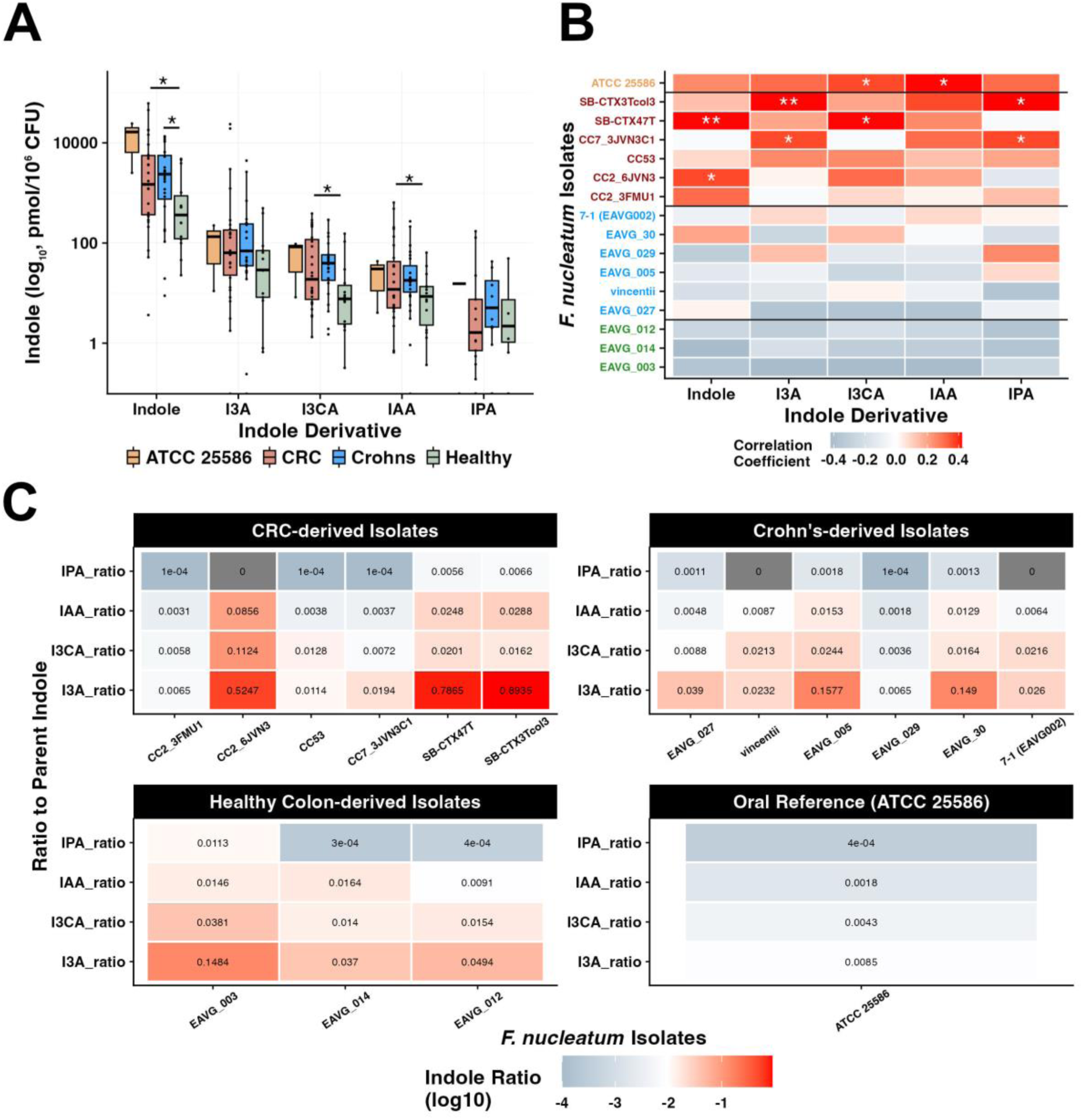
Isolates exhibit strain-specific indole derivative profiles and metabolic preferences. **(A)** HPLC quantification of indole and five derivatives in Columbia Broth spent media. Box plots illustrate the production of indole, I3A, I3CA, IAA, and IPA in pmol/10^6^ CFU. P-values indicate significant differences between clinical groups determined by one-way ANOVA. **(B)** Point-biserial correlation heatmap between individual F. nucleatum isolates and specific indole derivatives. The y-axis is color-coded by clinical origin: green (healthy colon), blue (Crohn’s), red (CRC), and orange (ATCC 25586). Asterisks denote statistically significant correlations (* *p* < 0.05; ** *p* < 0.01). **(C)** Ratios of indole derivatives to parent indole, indicating metabolic preference for specific pathways. Each cell displays the ratio value, with colors representing low (blue) to high (red) ratios. Isolates are grouped by clinical association: CRC (n=6), Crohn’s (n=6), healthy colon (n=3), and oral reference strain (ATCC 25586; n=1).

### *F. nucleatum* isolates demonstrate variability in biofilm formation across clinical origins

Prior to evaluating the effects of exogenous indole treatments, we first quantified baseline biofilm formation over 72 hours to determine if clinical origin inherently influenced *F. nucleatum* biomass accumulation. Biofilm development was monitored at 12-hour intervals over a 72-hour period. At the 12-hour mark, most isolates exhibited minimal biofilm development, except for the CRC-derived isolate CC2_3FMU1, which displayed the highest early-stage biomass **(Figure 3A)**. By 72 hours, distinct phenotypes emerged; the most robust biofilm formers were predominantly Crohn’s disease-derived (e.g., 7-1 (EAVG002) and EAVG_30), while most CRC-derived isolates, with the notable exception of SB-CTX3Tcol3, produced the lowest baseline biomass **(Figure 3A-B**; **Supplementary Figure 4)**. Consistent with our indole observations, these phenotypic distinctions diminished when isolates were grouped by subspecies (**Figure 3C**). Collectively, these data highlight substantial differences in biofilm formation across clinical isolates of *F. nucleatum*.

**Figure 3:**
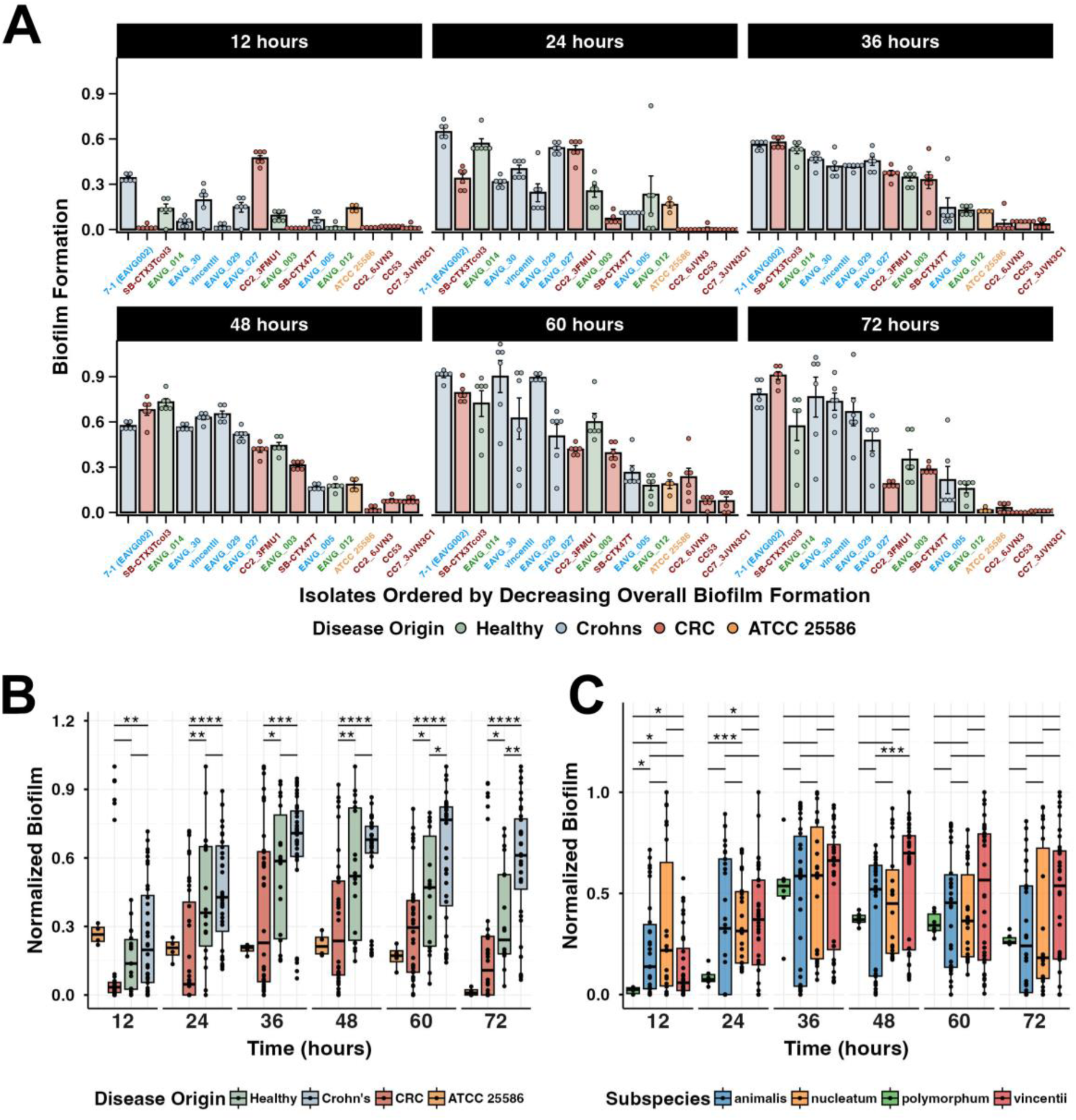
Variability in biofilm formation among clinical *F. nucleatum* isolates across different clinical origins. (A) Biofilm formation was quantified at six time points (12, 24, 36, 48, 60, and 72 hours) for 16 clinical isolates. Measurements were normalized to the maximum biofilm value at each time point. Statistical significance for differences between individual isolates was analyzed by adjusted ANOVA (Benjamini–Hochberg correction), followed by Tukey’s HSD test for pairwise comparisons. Significance is denoted as *p* < 0.05 (*), *p* < 0.01 (**), *p* < 0.001 (***), and *p* < 0.0001 (****). **(B, C)** Boxplots displaying normalized biofilm formation of isolates categorized by clinical origin **(B)** and subspecies **(C)**. Values are shown as the proportion relative to the maximum biofilm measurement at each time point. **Note:** Categorization by clinical origin includes CRC (n=6), Crohn’s (n=6), and healthy colon (n=3). The oral isolate (n=1) is included as a single reference point and was excluded from group-based statistical comparisons.

*F. nucleatum* exhibits isolate- and derivative-specific phenotypic shifts in response to exogenous indoles.

To ensure that subsequent virulence assays were not confounded by growth inhibition, we evaluated the impact of indole derivative exposure (0.25–2 mM) on isolate growth kinetics using area under the curve (AUC) analysis **(Supplementary Figure 6)**. While most isolates maintained stable growth up to 1 mM, some strains exhibited significant growth inhibition (e.g., CC53) or stimulatory phenotypes (e.g., CC2_FMU1). Similarly, indole derivatives modulated biofilm formation in a strain-specific manner **(Supplementary Figure 7)**.

To identify which *F. nucleatum* isolates were most sensitive to indole derivatives, we combined our growth and biofilm findings **(Supplementary Figure 5 and 6)** into a single metric: the maximum mean signed deviation **(Figure 4)**. By calculating the largest deviation from the untreated baseline for both growth and biofilm, we determined the magnitude and direction of each isolate’s general response to that indole. Hierarchical clustering of the max signed deviation distinguished specific response patterns among the isolates **(Figure 4A, Supplementary Figure 12)**. Notably, the CRC-derived isolate CC53 emerged as the most universally susceptible strain, exhibiting significant dual repression of both growth and biofilm across nearly all tested derivatives (p<0.001). In contrast, other sensitive strains like CC2_6JVN3 and ATCC 25586 displayed a decoupled stress response, characterized by severe growth attenuation paired with biofilm induction (p<0.05).

**Figure 4.**
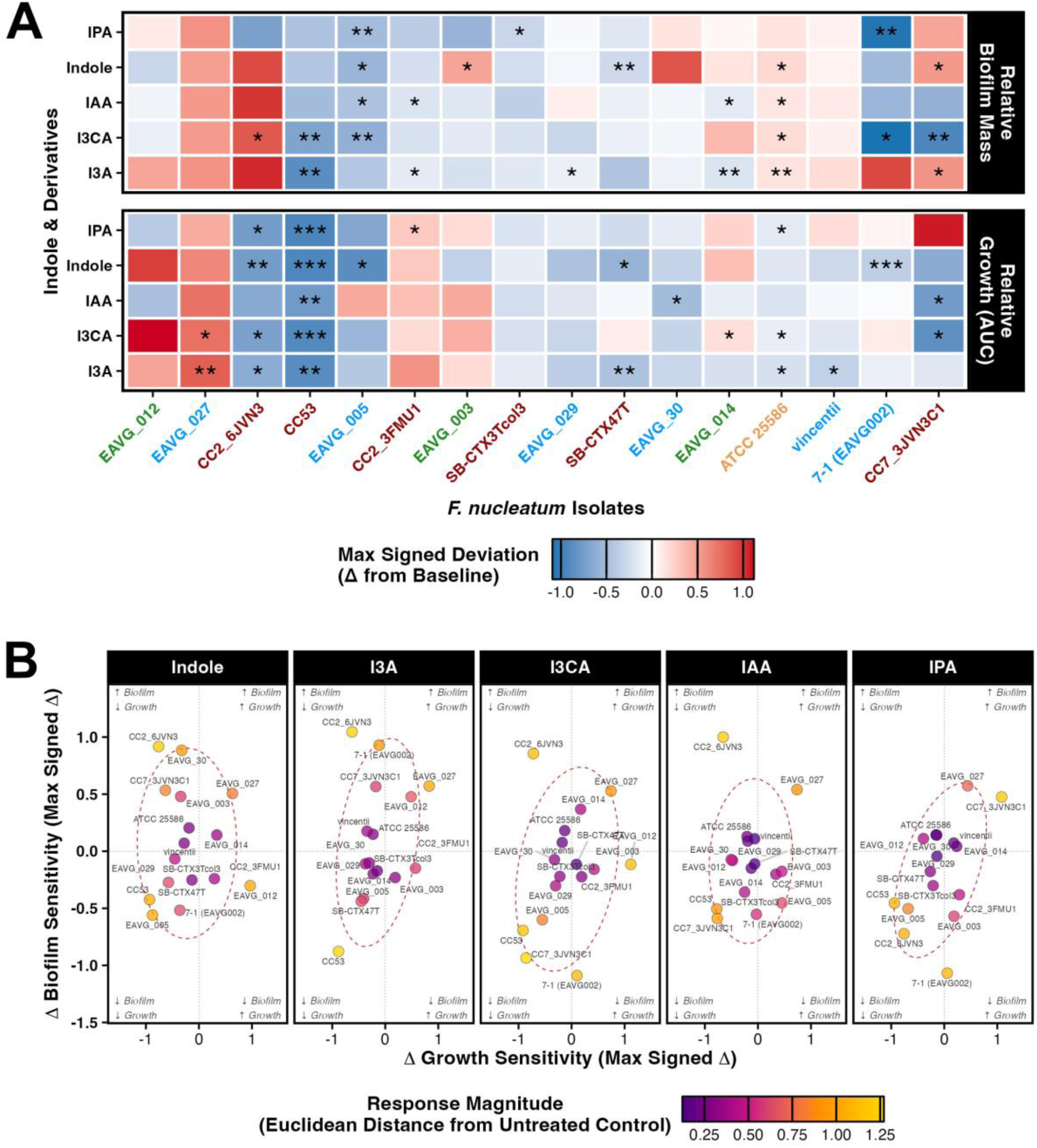
*Fusobacterium nucleatum* exhibits isolate- and derivative-specific phenotypic shifts in response to exogenous indoles. **(A)** Heatmap depicting the maximum signed deviation (calculated as the maximum mean difference (Δ) between the treatment and the untreated control across all tested concentrations) in relative biofilm mass and overall growth (AUC) for isolates exposed to indole and its derivatives. Phenotypic induction and repression are shown in red and blue, respectively. Isolates are hierarchically clustered (Ward’s method, Euclidean distance) and color-coded by clinical origin (red: CRC; blue: Crohn’s; green: Healthy colon; orange: Oral Reference). Significance versus untreated controls (* *p* < 0.05, ** *p* < 0.01, *** *p* < 0.001) was determined via one-way ANOVA with Benjamini-Hochberg FDR correction. **(B)** Bivariate scatter plots of coupled biofilm and growth maximum signed deviation responses per treatment. The origin (0,0) represents the untreated baseline. Point size and color denote the overall Response Magnitude, defined as the Euclidean distance from the baseline (√Δ*Growth*^2^ + Δ*Biofilm*^2^), highlighting hyper-responsive isolates in yellow.

To define which isolates were hyper-responsive, we mapped these coupled growth and biofilm changes and calculated each strain’s total response magnitude **(Figure 4B)**. This bivariate map separated the hyper-responsive isolates (CC53, EAVG_027, CC2_6JVN3, and 7-1 (EAVG002)), which consistently fell outside the 75% population confidence interval, the boundary representing the general response observed across the isolates. In contrast, indole-insensitive isolates clustered tightly around the origin (0,0), demonstrating their broad tolerance to indole-mediated stress across both growth and biofilm phenotypes. Interestingly, among hyper-responsive isolates, the nature of the sensitivity was highly specific to both the isolate and the specific indole derivative. While CC2_6JVN3 was universally responsive to every indole tested, other strains exhibited much more selective sensitivities; for instance, CC53 was predominantly affected by I3A, while 7-1 (EAVG002) was most sensitive to I3CA and IPA. These findings suggest that while most *F. nucleatum* isolates remain insensitive to exogenous indole derivatives, specific clinical isolates possess distinct vulnerabilities that are triggered by specific indole derivatives.

### Endogenous indole production is not a primary predictor of sensitivity to exogenous indole derivatives

We next asked whether an isolate’s endogenous indole production could predict its sensitivity to exogenous treatment **(Figure 5**). Comparison of total indole (Kovac’s reagent) with phenotypic responses revealed no significant correlation in either biofilm mass (ρ = 0.26, *p* = 0.338) or growth (ρ = 0.31, *p* = 0.239) **(Figure 5A)**. In fact, the isolates that produced the most indole (SB-CTX47T, SB-CTX3Tcol3) were largely insensitive to indole treatment except for the isolate CC7_3JVN3C1. The most sensitive isolates (EAVG_012, EAVG_027, 7-1 (EAVG002), CC53, EAVG005, and CC2_6JVN3) were low to moderate indole producers. To achieve higher resolution, we used HPLC to correlate the endogenous production of specific derivatives with their corresponding exogenous effects **(Figure 5B)**. Consistent with the Kovacs data, most indole derivatives exhibited a decoupled relationship between production and sensitivity. A notable exception occurred with I3CA, where endogenous levels showed a moderate negative correlation with biofilm response (ρ = −0.51, *p* = 0.048). This trend was largely driven by a subset of isolates (including 7-1 (EAVG002), CC7_3JVN3C1, CC53, and SB-CTX3Tcol3), which were both high I3CA producers and highly susceptible to I3CA-mediated biofilm inhibition. Together, these results suggest that *F. nucleatum* sensitivity to exogenous indoles are largely independent of its own indole production.

**Figure 5:**
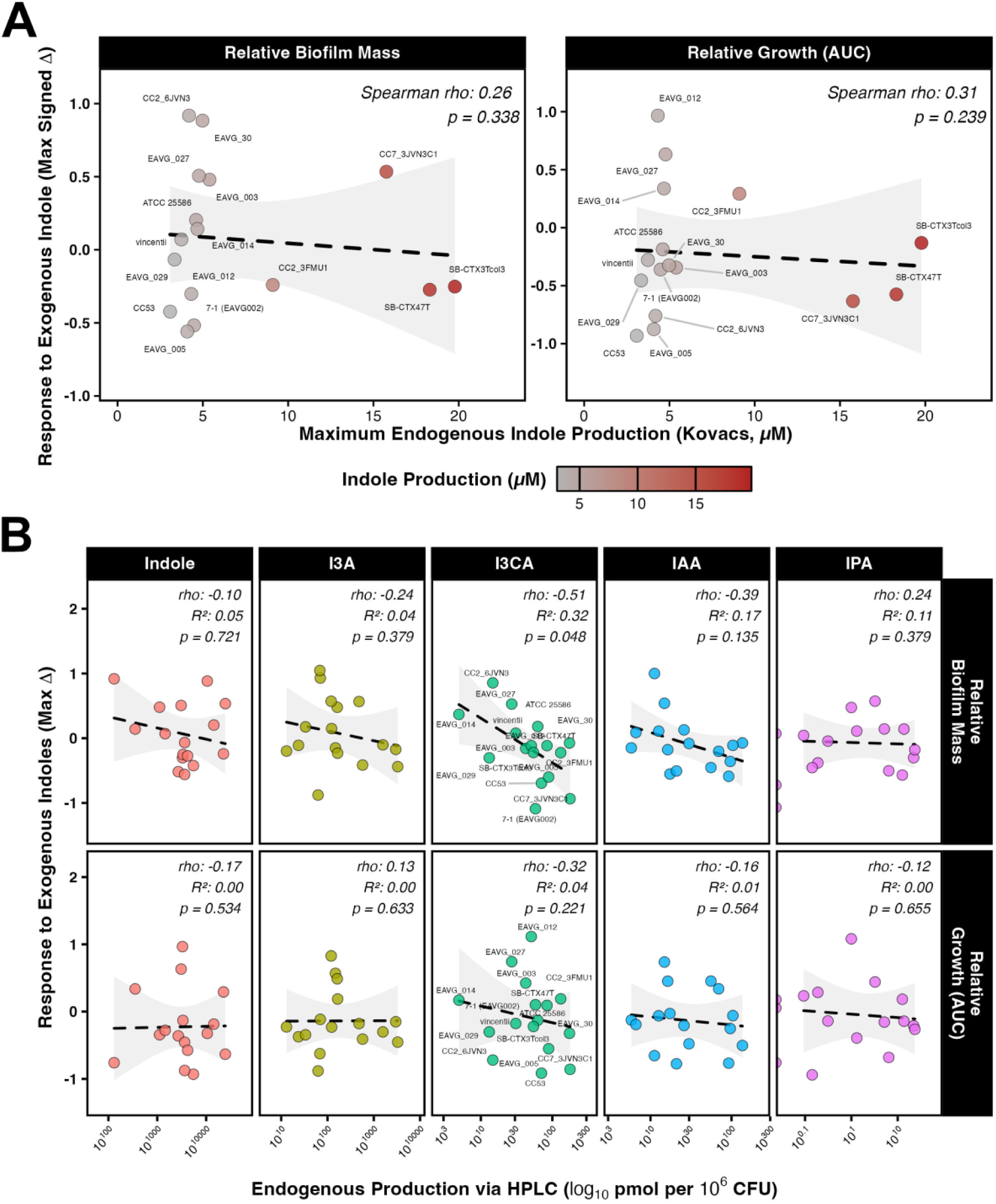
Endogenous indole production is independent of sensitivity to exogenous indoles. **(A)** Correlation between maximum endogenous indole production (quantified via Kovacs assay) and peak phenotypic response to exogenous indole (Max Signed Δ) across *F. nucleatum* isolates. Point fill scale corresponds to indole production concentration (µM). **(B)** Correlation between the endogenous production of specific indole derivatives (quantified via HPLC, log 10 pmol per 10^6^ CFU) and response to the corresponding exogenous treatment. For all plots, dashed lines represent linear regression trends with 95% confidence intervals (gray shading). Spearman correlation coefficients (*rho* = *ρ*), R^2^ values, and *p*-values are displayed.

### Transcriptional profiling reveals strain-specific heterogeneity and directional expression signatures

To determine if indole exposure induces distinct transcriptional responses, we quantified the expression of four virulence-associated genes (*aid1*, *fomA*, *fadA*, *radD*) and the tryptophanase gene (*tnaA*) across the 16 isolates following 72-hour exposure. Mapping these profiles onto a maximum likelihood cladogram revealed heterogeneous responses independent of genetic relatedness **(Figure 6A)**. Closely related isolates exhibited divergent expression (e.g., vincentii vs. CC53), while distantly related isolates showed convergent *fomA* upregulation. To evaluate whether these expression patterns correlate with clinical origin, we utilized a Naïve Bayes classification model. Certain genes, notably *fomA*, showed disease-specific expression signatures. These distinct frequency patterns **(Figure 6B)** provided the conditional probability framework for our Naïve Bayes classification model. The categorical expression direction model (Up vs. Down) yielded an AUC of 0.77 (*accuracy = 0.58, p = 0.026*), outperforming raw fold-change values **(Figure 6C, Supplementary Figure 13B)**. Precision-recall analysis further supported the stability of the directional model across cohorts **(Figure 6E, Supplementary Figure 14)**. Feature importance and univariate analyses identified *fomA*, *tnaA*, and *aid1* as the primary drivers of this association **(Figure 6D, Supplementary Figure 13D)**. While these data suggest a preliminary association between indole-induced transcriptional shifts and disease origin, we acknowledge the high risk of model overfitting given the limited sample size (n=16); consequently, these patterns represent exploratory, isolate-specific traits rather than definitive diagnostic markers. Moreover, clinical labels were scattered throughout the cladogram **(Figure 6A)**, showing that neither taxonomy nor disease origin dictates a uniform response. This disconnect suggests that indole sensitivity may be unique trait of individual isolates, rather than a shared evolutionary characteristic.

**Figure 6.**
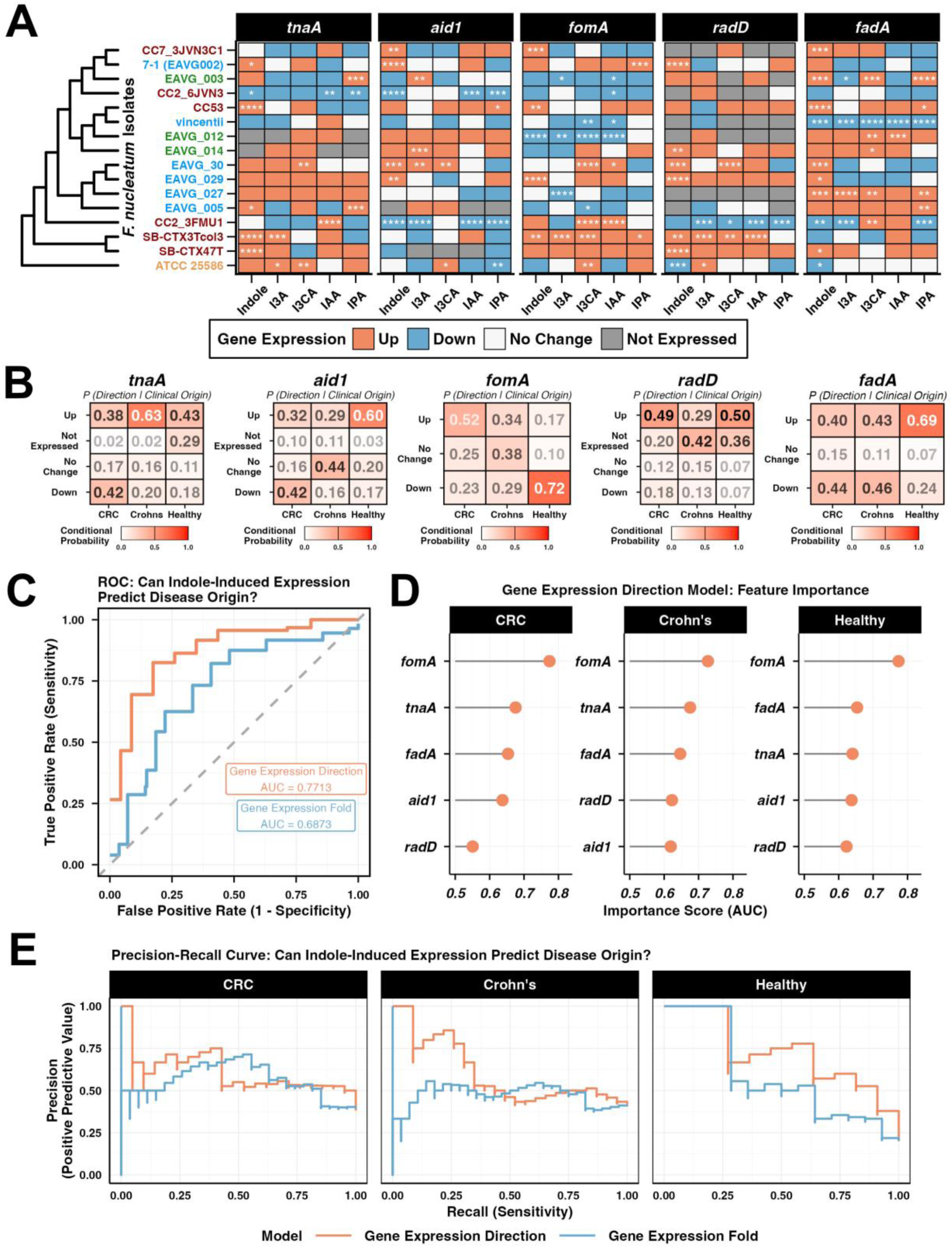
Transcriptional profiling reveals strain-specific heterogeneity and directional expression signatures. **(A)** 16S rRNA-based maximum likelihood cladogram of 16 *F. nucleatum* isolates paired with a heatmap showing expression of targeted virulence genes in response to 1 mM indole (72 h). Heatmap displays relative fold expression (2^-**ΔΔ**Ct^, normalized to 16S rRNA and vehicle control). Expression direction threshold is categorized as Up (fold > 1.25; red), Down (fold < 0.75; blue), or No Change (white; gray = undetected). Significance was determined via pairwise t-tests with Bonferroni correction (*p<0.05, **p<0.01, ***p<0.001, ****p<0.0001). **(B)** Conditional probabilities (*P (Direction | Clinical Origin)*) of gene expression states across clinical cohorts. These frequencies were derived from the heatmap in **(A)** and served as training features for the Naïve Bayes classifier. **(C)** Receiver Operating Characteristic (ROC) curve assessing Naïve Bayes performance (true positive vs. false positive rate) using categorical Expression Direction versus quantitative Fold Change. **(D)** Feature importance analysis of the categorical expression model. Scores show the AUC for individual genes, reflecting their independent capacity to classify a specific disease state against all others (0.5 = random prediction). **(E)** Precision-recall curves evaluating the trade-off between precision (positive predictive value) and recall (sensitivity) when classifying disease origin with the categorical versus quantitative models.

### Indole derivatives strengthen epithelial tight junction integrity and restrict bacterial invasion in CRC cell models

To determine if indole derivatives could influence virulence independently of growth and biofilm dynamics, we selected isolate SB-CTX3Tcol3 as this isolate remains phenotypically insensitive to treatment despite its high levels of endogenous indole production and biofilm formation. A bacterial invasion assay using Caco-2 cells demonstrated that indole treatment for 2 or 4 hours markedly inhibited invasion, producing a pathogen-suppressive effect comparable to the presence of antibiotics in the medium (**Figure 7A**). Notably, I3CA exhibited the strongest early inhibitory effect, reducing invasion by approximately 20% after only 2 hours. By 4 hours, treatment with indole, I3A, and I3CA suppressed invasion by more than 50%, whereas IAA and IPA had no detectable impact on the invasion capacity of SB-CTX3Tcol3. These data highlight that indole-induced effects are specific to the indole species present. The reliance of *F. nucleatum* on adhesins such as FadA to bind E-cadherin and promote invasion prompted us to investigate whether indoles could strengthen the intestinal barrier to prevent this process. When Caco-2 cells were treated with indole derivatives at the concentrations used in the invasion assay, transcripts for tight and adherens junction genes were predominantly upregulated (**Figure 7B**). Notably, I3A exerted the strongest effect on transcript levels, followed by indole, while I3CA, IAA, and IPA showed minimal impact on transcript expression. Similar, though less pronounced, trends were observed in HT-29 cells. These findings suggest that certain indoles enhance intestinal barrier integrity to help prevent pathogen invasion. However, the strong inhibitory effect of I3CA on bacterial entry despite its low impact on host gene transcripts suggests it may operate through a distinct, yet unidentified, mechanism.

**Figure 7:**
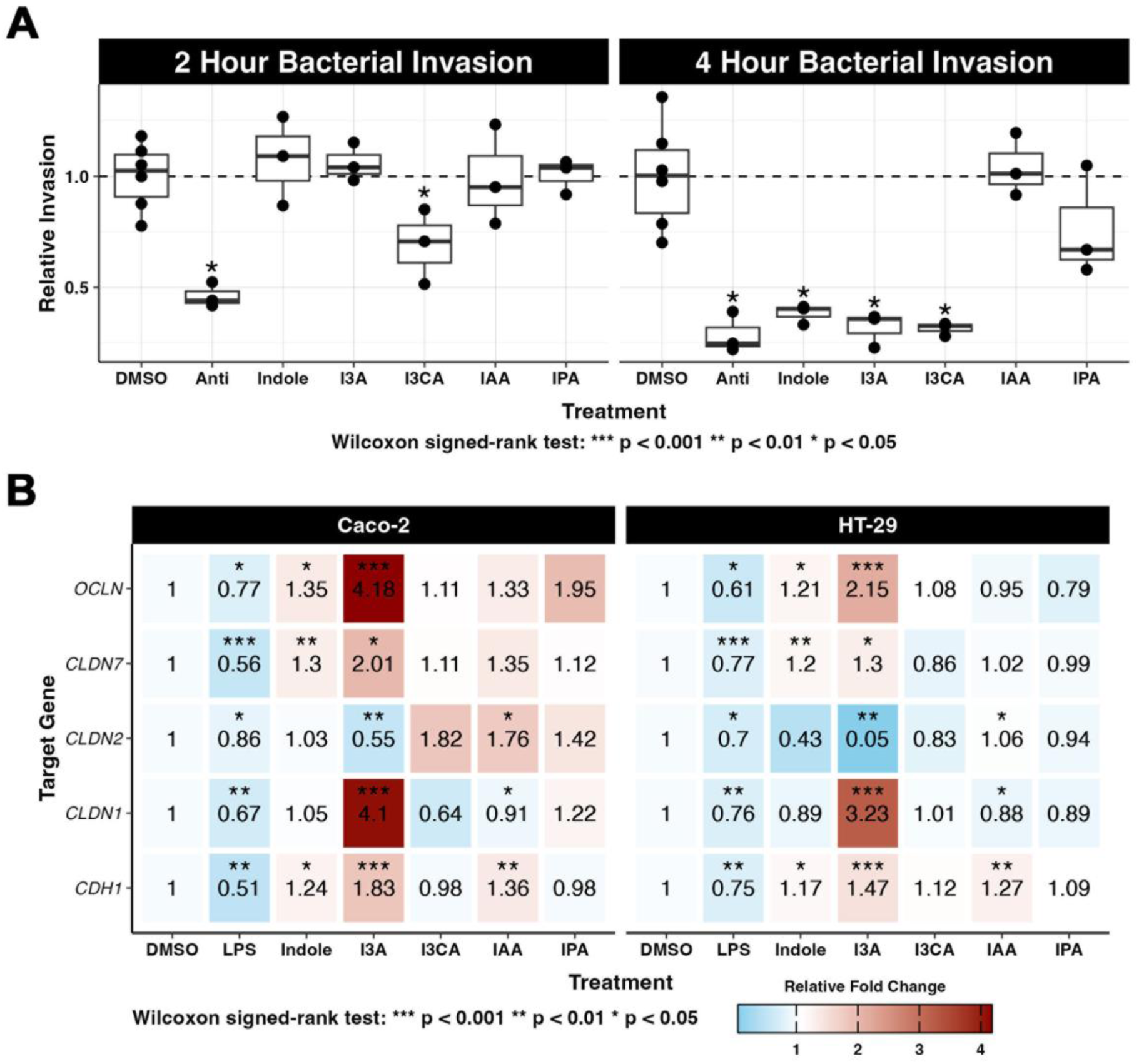
Indole derivatives strengthen epithelial tight junction integrity and restrict bacterial invasion in CRC cell models. **(A)** Invasion of a high-indole, high-biofilm producing CRC isolate (SB-CTX3Tcol3) into Caco-2 cells over 2 and 4 hours. *F. nucleatum* cultures were adjusted to a multiplicity of infection (MOI) of 50:1 and added to Caco-2 cells treated with 1 mM indole derivatives, vehicle control (0.1% DMSO), or a combination of penicillin (200 U/mL) and streptomycin (200 µg/mL) as a negative control (Anti). Statistical significance was assessed by Wilcoxon signed-rank test: ***p < 0.001, **p < 0.01, *p < 0.05. **(B)** Caco-2 and HT-29 cells were treated for 24 hours with 1 mM indole derivatives, vehicle control (0.1% DMSO), or lipopolysaccharide (LPS, 10 µg/mL) as a positive control for barrier disruption. Relative fold expression change (treatment vs. DMSO control) for tight junction genes (*CLDN1, CLDN2, CLDN7*) and adherens junction genes (*OCLN, CDH1*) was analyzed by qPCR and calculated using the *ΔΔCt* method. Heatmap colors represent the average fold change across four biological replicates, with pairwise comparisons assessed by Wilcoxon signed-rank test.

## DISCUSSION

In this pilot study, we investigated whether bacteria isolated from disease environments retain niche-specific virulence programs, particularly those governing biofilm formation and indole production. Using *F. nucleatum* as a model, we profiled indole production and sensitivity across 16 clinical isolates. Our central findings are that *F. nucleatum* clinical isolates exhibit significant strain-level heterogeneity across multiple phenotypic metrics, with diversity being attributed to individual isolates as opposed to disease origin or subspecies. Specifically, we observed : (i) substantial variation in indole production, with some isolates producing 3–4-fold more indole than average and a subset biased toward I3A derivative production; (ii) a wide range of biofilm formation capacities; (iii) isolate-dependent sensitivity to exogenous indole derivatives, including hyper-responsive clinical isolates; (iv) a relationship between increased I3CA production and I3CA-mediated biofilm inhibition; (v) highly heterogeneous, isolate-specific transcriptional responses to indole derivatives, where the direction of gene expression provides a stronger classification signal than expression magnitude; and (vi) indole derivative–specific strengthening of epithelial tight junctions while simultaneously reducing bacterial invasion. Together, these findings suggest that clinical isolates of *F. nucleatum* exhibit diverse indole metabolic programs that may influence virulence and adaptation within disease-associated environments.

Indole production is a defining feature of gut microbiota (37); however, the specific metabolic contributions of opportunistic pathogens like *F. nucleatum* within this network remain poorly defined. While recent evidence implicates tryptophan metabolites in inflammatory conditions and CRC pathogenesis (28,38,39), our data demonstrate that *F. nucleatum* is not a uniform participant in this process. Rather than exhibiting a conserved metabolic profile, we observed profound isolate-level heterogeneity in baseline indole production. Crucially, our isolates produced a diverse array of indole derivatives, most notably I3A. This metabolite was preferentially produced by a subset of CRC-derived isolates, consistent with previous reports identifying I3A as differentially abundant in CRC patient cohorts (36). Together, these findings demonstrate strain-specific metabolic diversity suggesting a functional consequence within the host.

Current understanding of *F. nucleatum* indole production is limited and disproportionately based on the ATCC 25586 reference strain (24,30,40). While IPA production in ATCC 25586 reportedly drives CRC progression via AhR-mediated M2 macrophage polarization (30), our data reveal that IPA is the least prevalent derivative across the clinical isolates examined, frequently falling below the limit of detection **(Figure 3A-C, Supplementary Figure 3A)**. This discrepancy suggests that IPA-driven procarcinogenic pathways may be isolate-specific rather than universal. By characterizing a broader range of clinical isolates, this work demonstrates that a single reference strain cannot capture the metabolic heterogeneity underlying indole’s role in *F. nucleatum* pathogenesis. These differences extend beyond indole production, as individual isolates also vary in how they respond to indole in their environment.

A defining characteristic of our study is the heterogeneity in *F. nucleatum* sensitivity to exogenous indole derivatives. While most responsive isolates (e.g., CC53) showed coordinated changes with growth and biofilm formation declining together, strains like CC2_6JVN3 and ATCC 25586 displayed a decoupled response, with biofilm formation increasing even as growth slowed. This pattern suggests that, for these isolates, indole may act as an environmental stressor, triggering a shift from planktonic growth to a protective biofilm state, a survival strategy observed in other bacterial pathogens (41). While most isolates exhibited only minor phenotypic variation, bivariate analysis of signed deviations highlighted three hyper-responsive strains: CC2_6JVN3, CC53, and 7-1 (EAVG002). CC2_6JVN3 showed broad sensitivity across nearly all derivatives, whereas CC53 and 7-1 displayed more selective vulnerabilities (7-1 primarily to I3CA and IPA, and CC53 to I3A). These sensitivities were largely independent of endogenous production, except for I3CA, where higher production correlated with greater biofilm inhibition (ρ = −0.51, p = 0.048). In contrast, robust indole producers like SB-CTX3Tcol3 remained largely insensitive, suggesting that these isolates may have evolved mechanisms to tolerate high local indole concentrations. Notably, the strongest phenotypic effects were observed at 2 mM, which may exceed typical bulk fecal indole concentrations; however, local concentrations within biofilms or the tumor microenvironment remain unknown and could be substantially higher (42). Together, these findings highlight that *F. nucleatum* strains display highly strain-specific sensitivity and insensitivity to indole derivatives.

Interestingly, the insensitivity to indoles appears to be decoupled from *F. nucleatum* invasion. At physiologically relevant concentrations (35,43), indole reduced invasion by ∼50% in the otherwise indole-insensitive and highest indole-producing isolate SB-CTX3Tcol3, indicating that indole can modulate invasive behavior independently of its antimicrobial or biofilm effects. These findings suggest that indole acts as a regulatory signal, suppressing active pathogenicity while simultaneously limiting population growth, as reflected in the reduced carrying capacity observed across most isolates **(Supplementary Figure 5)**. Given the high variability in both endogenous indole production and sensitivity, further mechanistic studies across a broader panel of clinical isolates are needed to definitively clarify indole’s role in *F. nucleatum* virulence.

Given our small sample size (n=16), any classification approach carries substantial overfitting risk. We therefore employed a Naïve Bayes classifier, leveraging its high-bias, low-variance architecture. The model achieved an AUC of 0.77; however, its modest accuracy (*0.58, p = 0.021 vs. NIR*) suggests it is better suited for exploratory pattern discovery than clinical diagnosis. Specifically, the analysis revealed that the direction of gene expression (up- vs. downregulation) predicted clinical origin more reliably than fold-change magnitude. This indicates that the core transcriptional logic of the adhesin *fomA* is highly conserved, even when the absolute strength of the response varies. As directional *fomA* expression was a primary classifier and was consistently upregulated in CRC-derived isolates, it may represent a viable target for strain-selective antimicrobials. Biologically, FomA serves as a critical anchor for host epithelial adherence and has demonstrated immune-adjuvant activity in mammalian models (16,17). This directional signal positions FomA as a compelling candidate for strain-selective therapeutic targeting, aligning with recent work demonstrating its viability as a target for precision antimicrobial peptide therapies (44). However, the broad dispersion of clinical origins across our cladogram confirms that these responses do not strictly track with phylogeny. Therefore, they should be considered exploratory signatures rather than definitive disease markers.

It is essential to acknowledge that this study serves as a pilot investigation. The limited number of isolates examined (n=16), the singular oral reference strain, and the inherent overfitting risks associated with modeling high-dimensional data on small cohorts mean that broader generalizations regarding disease or subspecies specificity require rigorous validation. Future research utilizing dual-RNA-seq during active invasion will be necessary to define the precise host-pathogen molecular pathways by which indoles impede bacterial entry. Despite these limitations, our findings establish that clinical isolates of *F. nucleatum* exhibit profound strain-level heterogeneity in indole metabolism and sensitivity. This metabolic diversity highlights vulnerabilities that represent candidate targets that can be exploited for precision or strain-selective therapeutic interventions.

## ACKNOWLEDGMENTS

The authors thank the Molecular Biosciences Core (MBC) at Baylor University for access to instrumentation and expert technical assistance. An earlier pre-print version of this manuscript is available on bioRxiv (45). We would also like to thank Dr. Emma Allen Vercoe and Dr. Susan Bullman for gifting the clinical *F. nucleatum* isolates.

## CRediT AUTHORSHIP CONTRIBUTION STATEMENT

**C.S.** Conceptualization, Data curation, Formal analysis, Investigation, Methodology, Project administration, Software, Validation, Visualization, Writing - original draft, Writing - review & editing. **A.C.** Conceptualization, Formal analysis, Investigation, Methodology, Software, Validation, Visualization, Writing – review & editing. **M.R.** Investigation, Methodology, Writing – review & editing. **J.H.** Data curation, Investigation, Writing – review & editing. **L.C.** Investigation, Writing – review & editing. **G.Z.** Investigation, Writing – review & editing. **R.L.** Conceptualization, Resources, Writing – review & editing. **L.G.** Corresponding author, Conceptualization, Funding acquisition, Methodology, Resources, Supervision, Writing – original draft, Writing – review & editing.

## DATA AVAILABILITY STATEMENT

The authors confirm that the data supporting the findings of this study are available within the article and its supplementary materials. All data generated or analyzed during this study have been deposited in the Texas Data Repository (https://doi.org/10.18738/T8/3PYJA9) and are available upon request.

---

**Table 1.** Sources and references of clinical Fusobacterium nucleatum isolates. A summary of clinical *F. nucleatum* isolates obtained from various disease contexts, including healthy colon controls, Crohn’s disease (remission and active), colorectal cancer, and oral lesions. Isolates are categorized by strain, alternative names (based on PubMed identifiers), sampling site, associated disease state, and subspecies classification. Subspecies was determined with subspecies specific primers (Supplemental Table 1). Supplier information and relevant literature references are also provided.

| STRAIN | ALT NAMES<br>PMID#<br>25370491 | SAMPLING SITE | DISEASE | SUBSPECIES | ISOLATE SUPPLIERS |
| --- | --- | --- | --- | --- | --- |
| *F. nucleatum animalis* EAVG_003 | 11_3_2 | Nonspecific colon biopsy | Healthy colon | *animalis* | Dr. Emma Allen-Vercoe, University of Guelph |
| *F. nucleatum* EAVG_014 | 3_1_27 | Rectum biopsy | Healthy colon | *vincentii* | Dr. Emma Allen-Vercoe, University of Guelph |
| *F. nucleatum* EAVG_012 | 4_1_13 | Ascending colon biopsy | Healthy colon | *vincentii* | Dr. Emma Allen-Vercoe, University of Guelph |
| *F. nucleatum* EAVG_027 | 2_1_50A | Distal colon biopsy | Crohn's remission | *animalis* | Dr. Emma Allen-Vercoe, University of Guelph |
| *F. nucleatum* subsp. *vincentii* | 1_1_2 | Distal colon biopsy | Crohn's remission | *vincentii* | Dr. Emma Allen-Vercoe, University of Guelph |
| *F. nucleatum* EAVG_005 | 13_3C | Sigmoid colon biopsy | Crohn's remission | *nucleatum* | Dr. Emma Allen-Vercoe, University of Guelph |
| *F. nucleatum* EAVG_029 | 4_1_31B | Ascending colon biopsy | Crohn's active | *vincentii* | Dr. Emma Allen-Vercoe, University of Guelph |
| *F. nucleatum* EAVG_030 | 3_1_36B3 | Ascending colon biopsy | Crohn's active | *vincentii* | Dr. Emma Allen-Vercoe, University of Guelph |
| *F. nucleatum* subsp. *nucleatum* CC2_3FMU1 |  | Tumor biopsy | Colorectal cancer | *nucleatum* | Dr. Emma Allen-Vercoe, University of Guelph |
| *F. nucleatum* subsp. *animalis* CC2_6JVN3 |  | Tumor biopsy | Colorectal cancer | *animalis* | Dr. Emma Allen-Vercoe, University of Guelph |
| *F. nucleatum* subsp. *vincentii* CC53 |  | Tumor biopsy | Colorectal cancer | *vincentii* | Dr. Emma Allen-Vercoe, University of Guelph |
| *F. nucleatum* subsp. *animalis* CC7_3JVN3C1 |  | Tumor biopsy | Colorectal cancer | *animalis* | Dr. Emma Allen-Vercoe, University of Guelph |
| *F. nucleatum* (SB-CTX47T) |  | Tumor biopsy | Colorectal cancer | *polymorphum* | Dr. Susan Bullman, MD Anderson Cancer Center |
| *F. nucleatum* (SB-CTX3Tcol3) |  | Tumor biopsy | Colorectal cancer | *nucleatum* | Dr. Susan Bullman, MD Anderson Cancer Center |
| *F. nucleatum* EAVG002-7_1-HM_73 | 7_1 | Ascending colon biopsy | Crohn's active | *animalis* | Dr. Emma Allen-Vercoe, University of Guelph |
| *F. nucleatum* subsp. *nucleatum* Knorr 25586 |  | ATCC | Oral Lesion | *nucleatum* | ATCC |

