## Supplementary Figures for "Clinical isolates of *Fusobacterium nucleatum* display strain-specific virulence and modulation by indole derivatives"

| Table 1: *F. nucleatum* Virulence Factor and Subspecies Primers | | | |
|---|---|---|---|
| **PRIMER** | **SEQUENCE** | **TARGET** | **REFERENCES** |
| FadA FWD | GCTAGACAAGCACTAGCACAA | Fusobacterium adhesin A (FadA) | In house |
| FadA REV | CTTGTGTTAAGCTTCTGCTTGAAGT | | |
| tnaA FWD | CGTGCATTAAAAGCAACTCACTATAATG | Tryptophanase enzyme | |
| tnaA REV | CAGTAGTTCCCGAGTCTGATAAC | | |
| aid1 FWD | TACAGGAGGTGCCGTAGCAG | aid1 (Adherence Inducing Determinant 1) | (1) |
| aid1 REV | TTTTTGTTAATTCTCCAGCTCCA | | |
| 16s FWD | TTGGACAATGGACCGAGAGT | F. nucleatum 16S rRNA | |
| 16s REV | GCCGTCACTTCTTCTGTTGG | | |
| RadD FWD | GGATTTATCTTTGCTAATTGGGGAAATTATAG | Fn1526 (now designated as radD) | |
| RadD REV | ACTATTCCATATTCTCCATAATATTTCCCATTAGA | | |
| FomA FWD | AGAGTTTGATCCTGGCTCAG | outer membrane protein porin (FomA) | |
| FomA REV | GTCATCGTGCACACAGAATTGCTG | | |
| Subsp. animalis FWD | TTTGTATTTGACAATTTAATTAGAGCA | Subspecies-specific PCR: Fn. animalis hypothetical protein gene CDS | (2) |
| Subsp. animalis REV | TGGGATAATAATCCAAATATYGCAC | | |
| Subsp. nucleatum FWD | CTTCATTGTTAGGATTACAAGAATAGGC | Subspecies-specific PCR: Fn. nucleatum Ankyrin repeat protein gene CDS | |
| Subsp. nucleatum REV | TTTCAGTAACATTTATTGGGAGGTTT | | |
| Subsp. polymorphum FWD | AAGATTAAATAAAGATGGAAATAACCATRAA | Subspecies-specific PCR: Fn. polymorphum hypothetical protein gene CDS | |
| Subsp. polymorphum REV | CCATCTCCAAAGCCTTTCC | | |
| Subsp. vincentii FWD | CTTTGTCTTATGATTTATTGCRTATCTTC | Subspecies-specific PCR: Fn. vincentii hypothetical protein gene CDS | |
| Subsp. vincentii REV | GAAGAATCAGAAGCACTACCAGCA | | |
| ACTB FWD | GGATCAGCAAGCAGGAGTATG | Housekeeping gene PCR: β-actin (ACTB) | In house |
| ACTB REV | AGAAAGGGTGTAACGCAACTAA | | |
| CDH1 FWD | CTCGACACCCGATTCAAAGT | E-cadherin (CDH1) | In house |
| CDH1 REV | CCAGGCGTAGACCAAGAAAT | | |
| OCLN FWD | CCCATCTGACTATGTGGAAAGAG | Occludin (OCLN) | In house |
| OCLN REV | AACCGGCGTGGATTTATAGG | | |
| CLDN 1 FWD | CCGTTGGCATGAAGTGTATGA | Claudin-1 (CLDN1) | In house |
| CLDN 1 REV | GCCAGACCTGCAAGAAGAAATA | | |
| CLDN 2 FWD | GATCCTACGGGACTTCTACTCA | Claudin-2 (CLDN2) | In house |
| CLDN 2 REV | CAGGGAGAACAGGGAAGAAATAA | | |
| CLDN 7 FWD | GGGAGACGACAAAGTGAAGAAG | Claudin-7 (CLDN7) | In house |
| CLDN 7 REV | ATACCAGGAGCAAGCTACCA | | |

**Supplementary Table 1: Primer sequences targeting *F. nucleatum* virulence factors, tight and adheren junction genes, and subspecies-specific genes.** Forward and reverse primer sequences used for qPCR amplification of key *F. nucleatum* virulence genes (e.g., fadA, aid1) and subspecies specific markers, along with their corresponding targets and references. References: 1.) Kaplan A, Kaplan CW, He X, McHardy I, Shi W, Lux R. Characterization of aid1, a novel gene involved in Fusobacterium nucleatum interspecies interactions. Microb Ecol. 2014 Aug;68(2):379–87. 2.) Krieger M,

AbdelRahman YM, Choi D, Palmer EA, Yoo A, McGuire S, et al. Stratification of Fusobacterium nucleatum by local health status in the oral cavity defines its subspecies disease association. Cell Host & Microbe. 2024 Apr 10;32(4):479-488.e4.

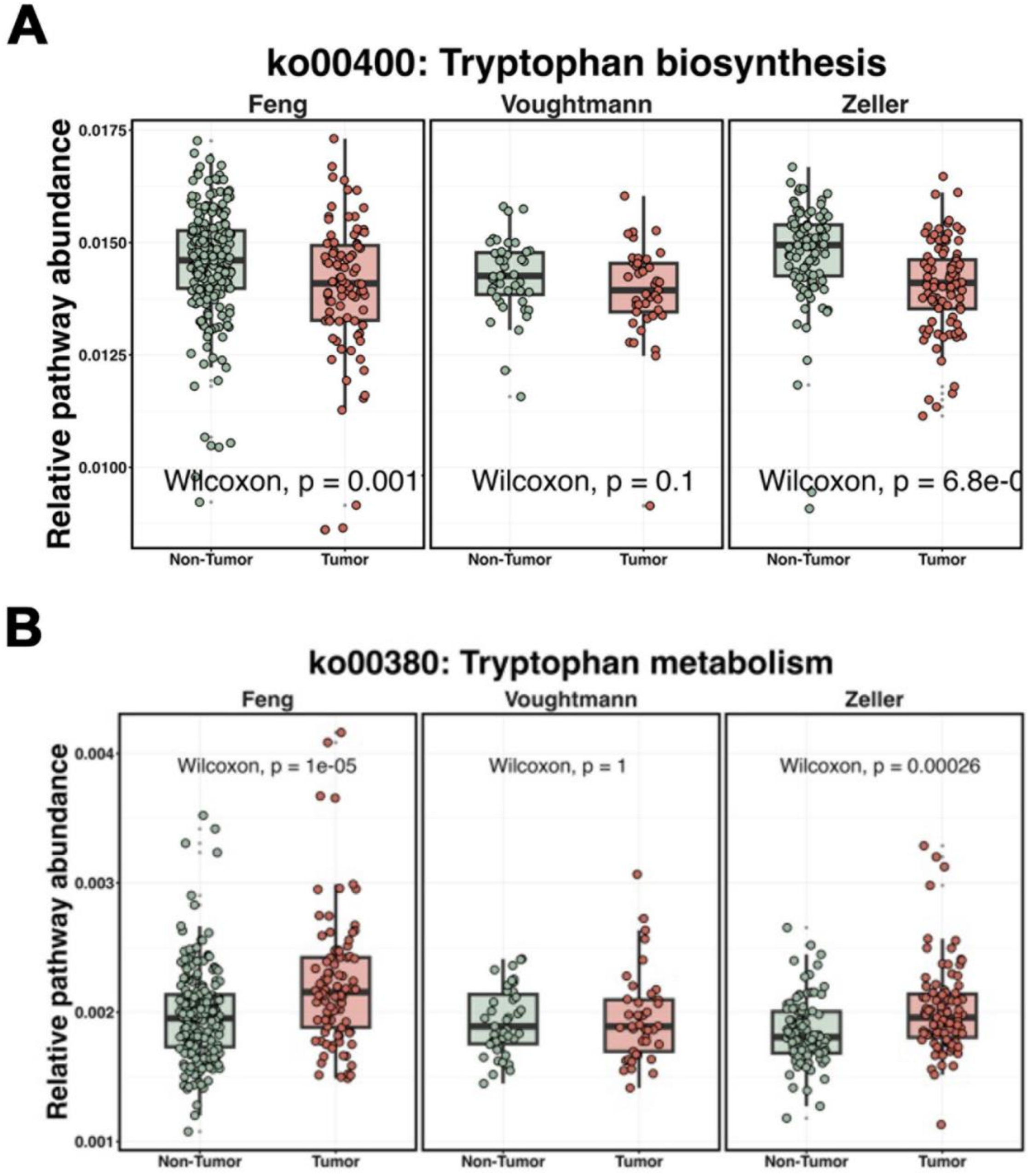


**Supplementary Figure 1: Meta-analysis of tryptophan metabolism and biosynthesis is significantly different in CRC vs health controls**. Relative pathway abundance inferred from gene markers analysis (PICRUSt) of fecal microbiome sequencing for tryptophan metabolism **(A)** and biosynthesis **(B)**. Represented data was reanalyzed from Feng 2015, Voughtmann 2015, and Zeller 2014.

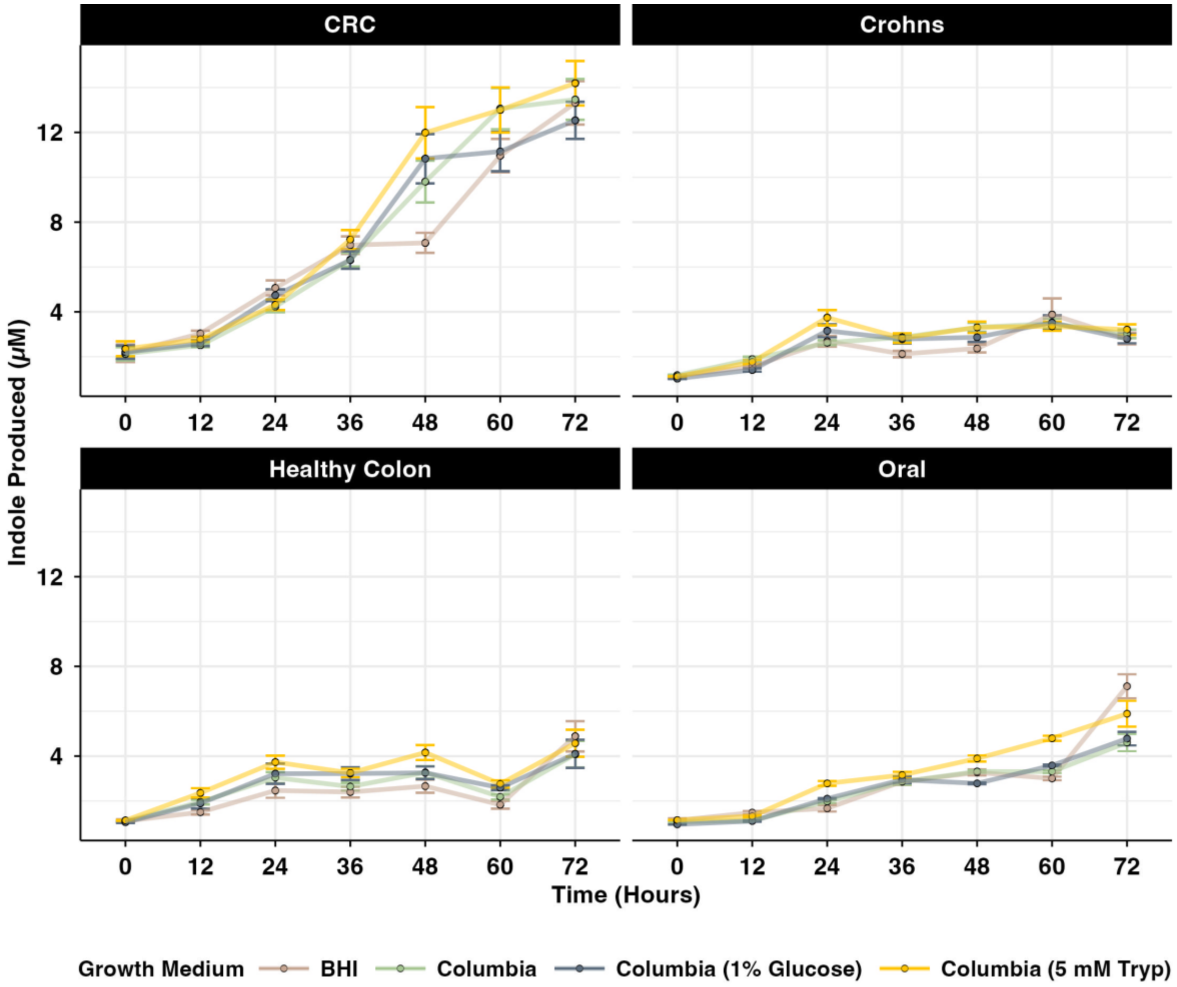


**Supplementary Figure 2: Impact of growth media on total indole production in *F. nucleatum* isolates.** * Bacterial isolates (n=16) were cultured over 72 hours in four distinct media: brain heart infusion (BHI), CB, CB + 1% glucose, and Columbia + 5 mM tryptophan (Tryp). Total indole was quantified at 12-hour intervals using Kovac's reagent and calibrated against a synthetic indole standard curve (0-2000μM). * Data are presented in Panel (A) as mean indole concentration (μM) +/- SEM, stratified by isolate disease origin: CRC (n=6), Crohn's (n=6), healthy colon (n=3), and oral (n=1).

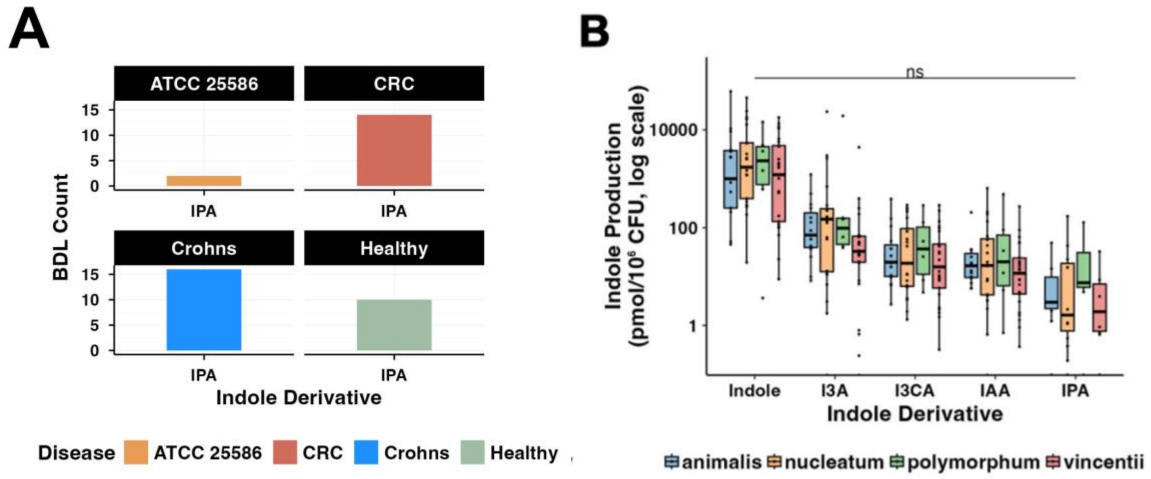


**Supplementary Figure 3: Analysis of indole derivative production across *Fusobacterium* subspecies. (A)** Distribution of indole-3-propionic acid (IPA) measurements falling below the analytical detection limit (BDL count) of the HPLC system across disease groups. **(B)** Concentration of five targeted indole derivatives—indole, indole-3-aldehyde (I3A), indole-3-carboxylic acid (I3CA), indole-3-acetic acid (IAA), and IPA—produced by isolates grouped by subspecies: *animalis* (n=5), *nucleatum* (n=4), *polymorphum* (n=1), and *vincentii* (n=6). Derivatives were quantified from spent media via HPLC with native fluorescence detection and normalized to bacterial density (pmol per $10^6$ CFU).  are presented as box plots representing the median, interquartile range (IQR), and 1.5 × IQR whiskers. Differences between subspecies were evaluated by one-way ANOVA; "ns" indicates no significant difference ($p > 0.05$).

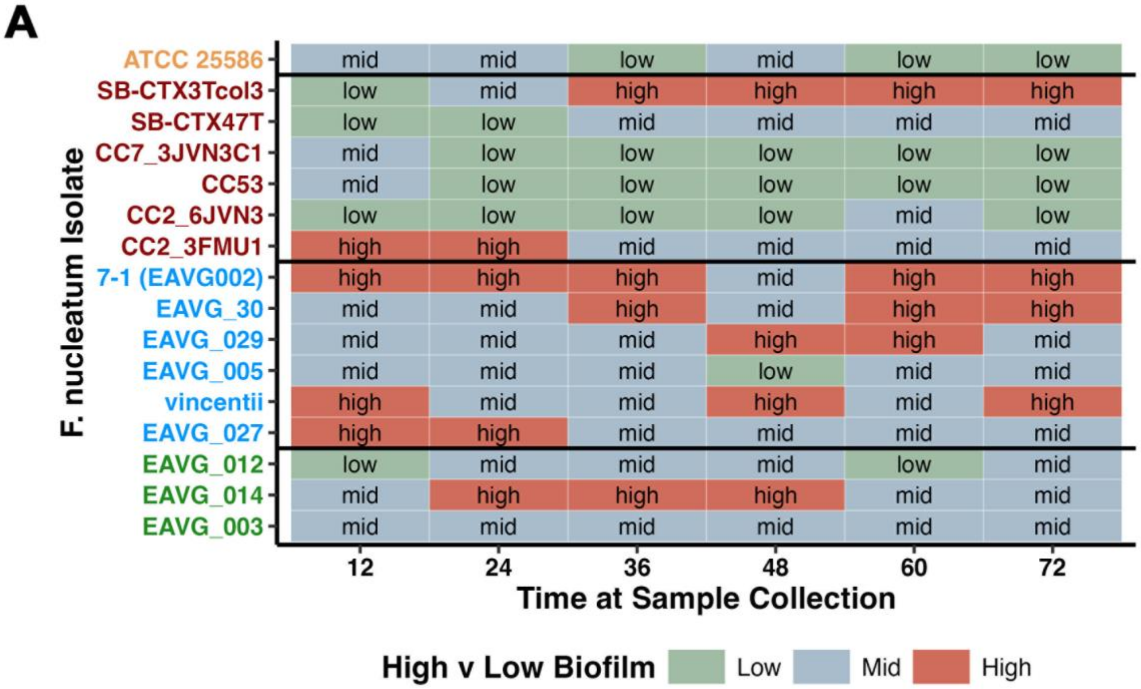


| F. nucleatum Isolate | 12 | 24 | 36 | 48 | 60 | 72 |
|---|---|---|---|---|---|---|
| ATCC 25586 | mid | mid | low | mid | low | low |
| SB-CTX3Tcol3 | low | mid | high | high | high | high |
| SB-CTX47T | low | low | mid | mid | mid | mid |
| CC7_3JVN3C1 | mid | low | low | low | low | low |
| CC53 | mid | low | low | low | low | low |
| CC2_6JVN3 | low | low | low | low | mid | low |
| CC2_3FMU1 | high | high | mid | mid | mid | mid |
| 7-1 (EAVG002) | high | high | high | mid | high | high |
| EAVG_30 | mid | mid | high | mid | high | high |
| EAVG_029 | mid | mid | mid | high | high | mid |
| EAVG_005 | mid | mid | mid | low | mid | mid |
| vincentii | high | mid | mid | high | mid | high |
| EAVG_027 | high | high | mid | mid | mid | mid |
| EAVG_012 | low | mid | mid | mid | low | mid |
| EAVG_014 | mid | high | high | high | mid | mid |
| EAVG_003 | mid | mid | mid | mid | mid | mid |

**Supplementary Figure 4: Dynamic biofilm production profiles of *F. nucleatum* isolates. (A)** Heatmap classifying isolates as *low*, *mid*, or *high* biofilm formers based on mean production levels relative to the cohort at that time. Thresholds were defined by quartiles (<25$^{th}$, 25$^{th}$-75$^{th}$ and >75$^{th}$ percentiles, respectively). Y-axis labels are color by disease association. Data represent measurements taken at 12 h intervals over a 72-h period.

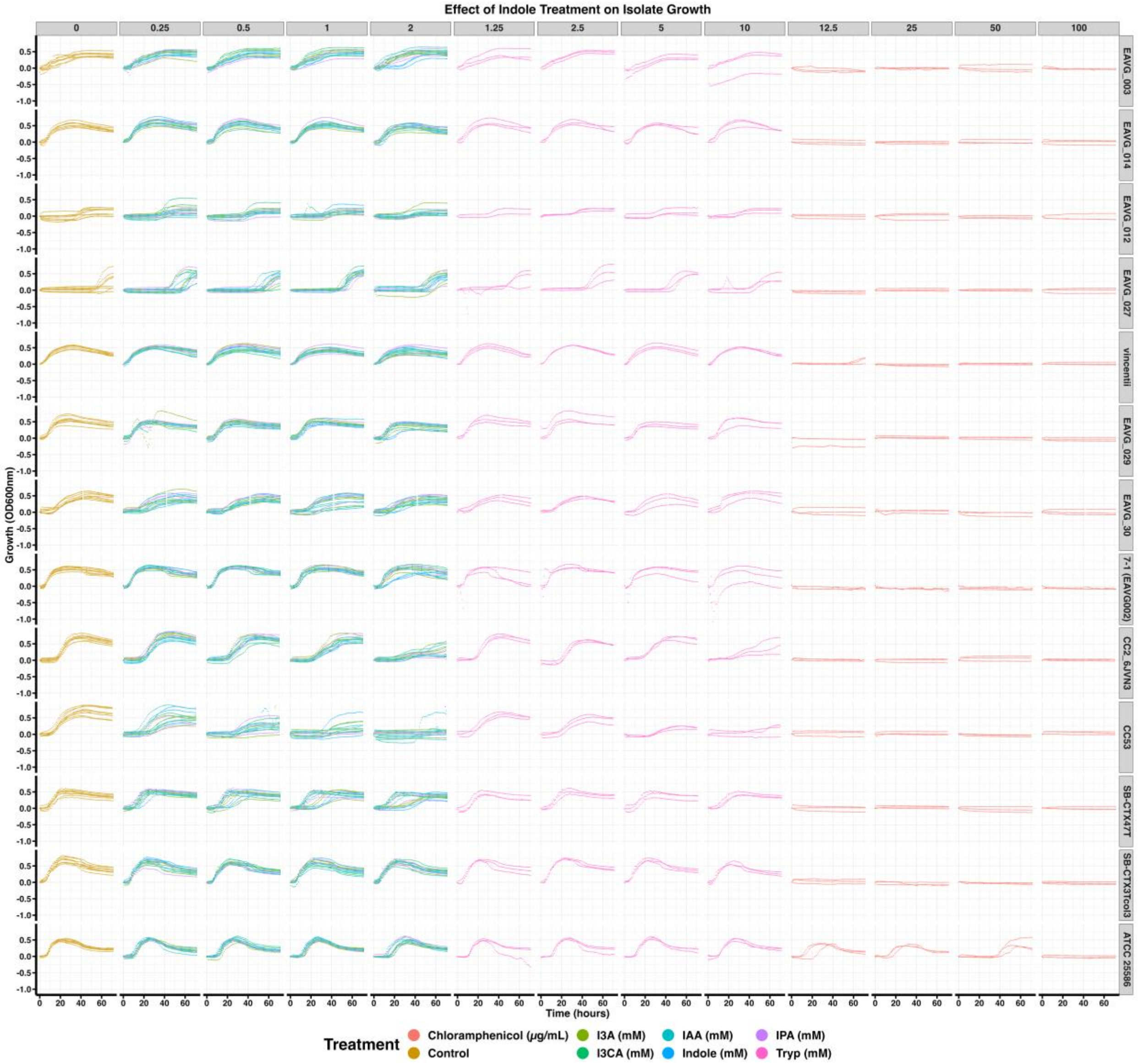


**Supplementary Figure 5: Concentration-dependent impact of indole derivatives on *F. nucleatum* growth kinetics.** Growth dynamics ($OD_{600}$) of 16 clinical isolates were monitored anaerobically over 72 hours. Treatment Gradient: Isolates were exposed to a concentration gradient (0, 0.25, 0.5, 1, and 2 mM) of five indole derivatives: indole, indole-3-aldehyde (I3A), indole-3-acetic acid (IAA), indole-3-propionic acid (IPA), and indole-3-carboxylic acid (I3CA).Tryptophan (Tryp) was included to assess metabolic conversion effects, and chloramphenicol (12.5–100 µg/mL) served as a positive control for growth inhibition. Each subplot displays growth curves for a single treatment concentration across all isolates to highlight inter-strain heterogeneity. Color-coded lines represent distinct treatments; data points reflect the mean of three biological replicates.

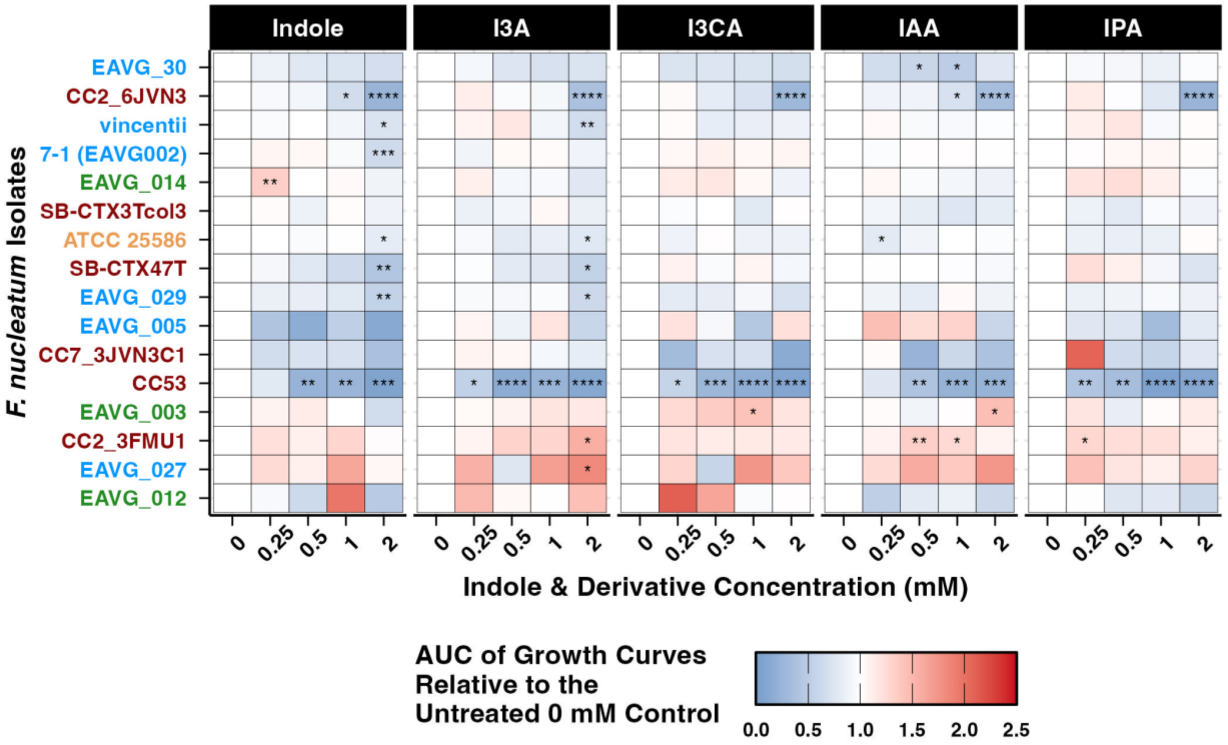


**Supplementary Figure 6: *Fusobacterium nucleatum* clinical isolates display strain-specific growth tolerances to exogenous indole and its derivatives.** Heatmap of Area Under the Curve (AUC, calculated with trapezoidal rule) for isolate growth over 72 hours exposed to indole, I3A, I3CA, IAA, and IPA, normalized to respective untreated (0 mM) controls. Colors indicate relative growth stimulation (red, >1) or inhibition (blue, <1). Isolates are ordered via hierarchical clustering of overall growth profiles using Euclidean distance and complete linkage. Asterisks denote significance versus 0 mM controls (*p < 0.05, **p < 0.01, ***p < 0.001, ****p < 0.0001; one-way ANOVA with Tukey's HSD, n=9 biological replicates for controls, n=3 biological replicates per treatment condition). Y-axis text colors represent isolate's disease origin (CRC-derived= red, Crohn's-derived=blue, Healthy colon-derived=green, and oral reference isolate=orange).

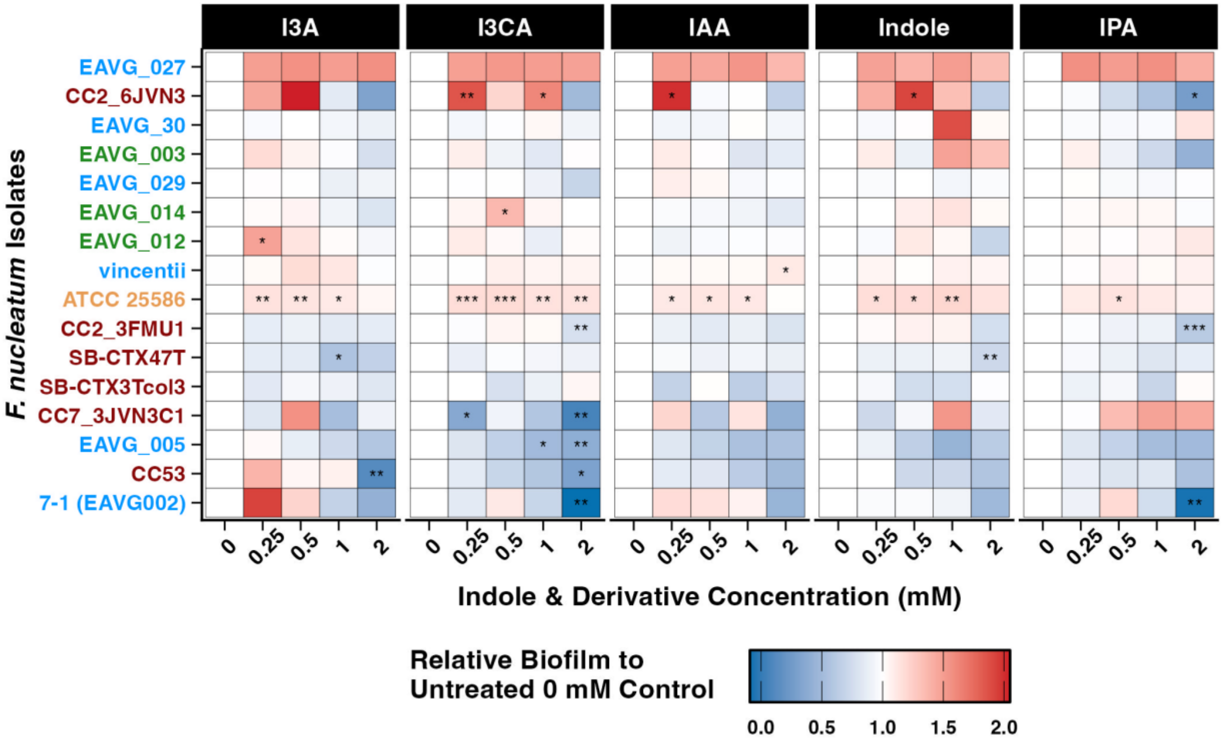


**Supplementary Figure 7. Impact of indole and its derivatives on *F. nucleatum* biofilm formation.** Heatmap displaying the relative biofilm production of isolates following treatment with varying concentrations (0, 0.25, 0.5, 1, and 2 mM) of I3A, I3CA, IAA, indole, and IPA. Biofilm biomass is expressed relative to the untreated control (0 mM) for each respective isolate. Colors indicate relative growth stimulation (red, >1) or inhibition (blue, <1). Isolates are ordered via hierarchical clustering of overall growth profiles using Euclidean distance and complete linkage. Asterisks denote significance versus 0 mM controls (*p < 0.05, **p < 0.01, ***p < 0.001, ****p < 0.0001; one-way ANOVA with Tukey's HSD, n=9 biological replicates for controls, n=3 biological replicates per treatment condition). Y-axis text colors represent isolate's disease origin (CRC-derived= red, Crohn's-derived=blue, Healthy colon-derived=green, and oral reference isolate=orange).

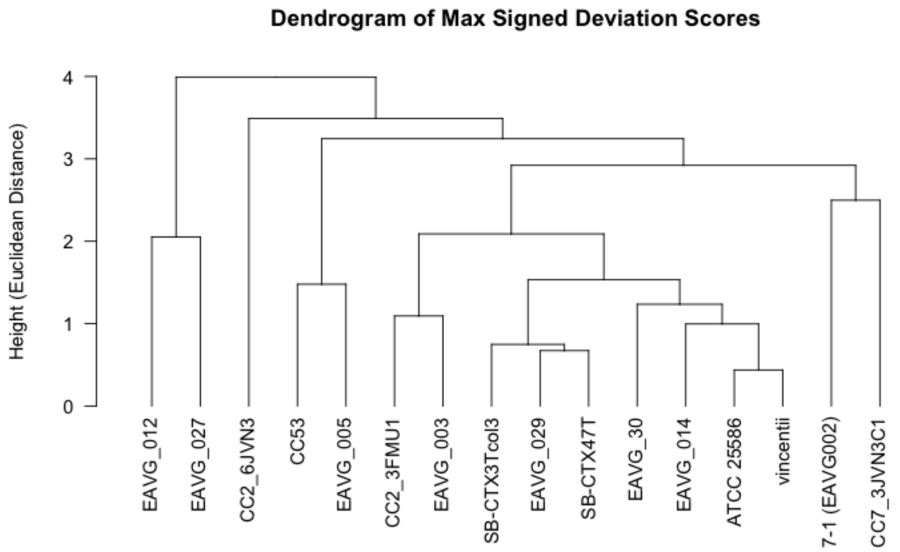


**Supplementary Figure 8: Hierarchical clustering of bacterial isolates based on indole sensitivity profiles.** Dendrogram representing the phenotypic similarity between 16 isolates based on maximum signed deviation scores across all indole derivatives and assays. Distances were calculated using Euclidean distance and clustered via Ward's D2 method. Vertical height represents the degree of divergence in response profiles, with lower heights indicating greater similarity in sensitivity patterns.

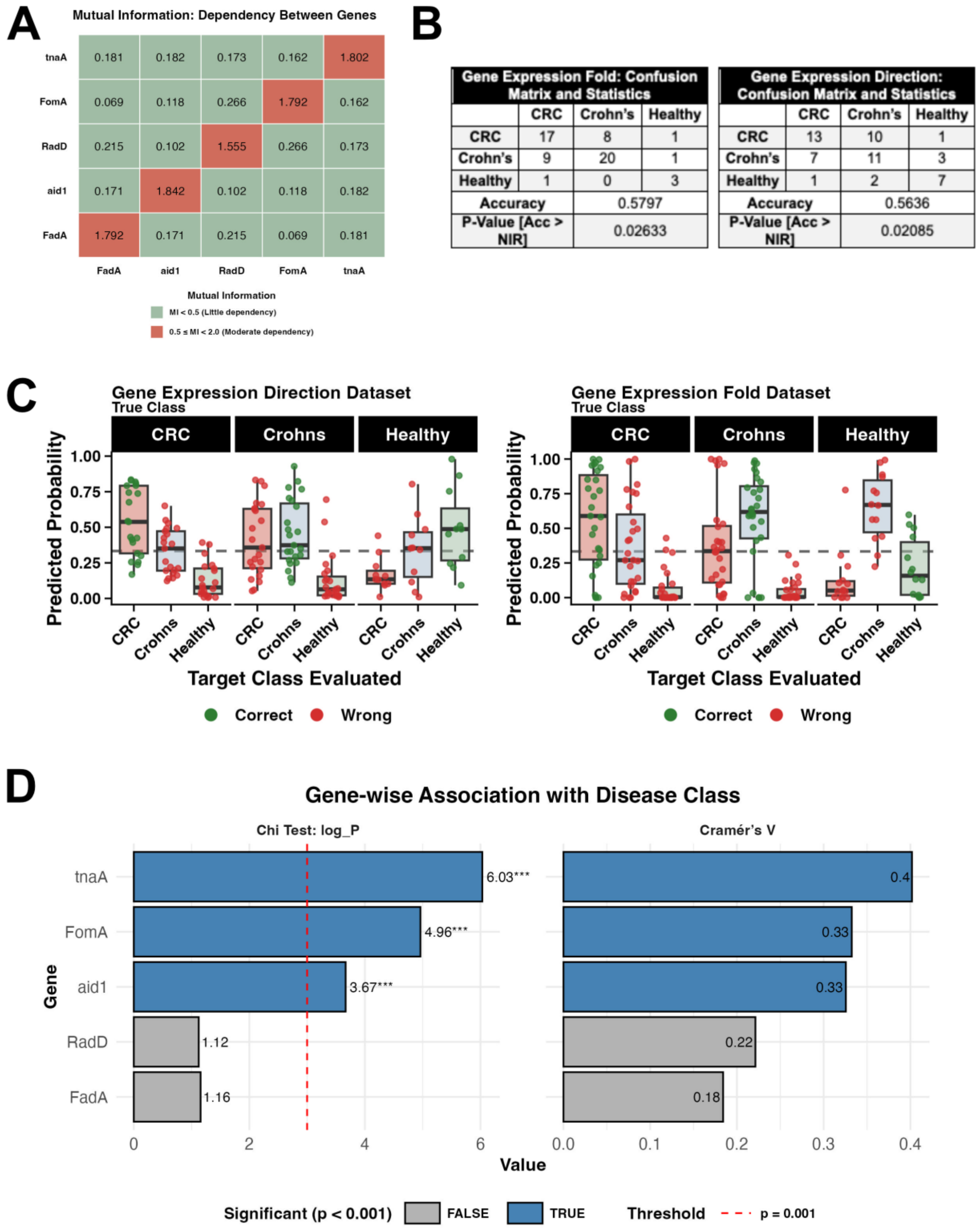


**Supplementary Figure 9: Statistical validation and performance metrics of the Naïve Bayes classification models. (A)** Mutual Information (MI) matrix assessing feature independence among indole-induced virulence and metabolic genes. Green tiles indicate low dependency (MI < 0.5), and red tiles indicate moderate dependency (0.5 ≤ MI < 2.0). **(B)** Confusion matrices and summary statistics, including overall accuracy and No Information Rate (NIR) p-values, for the quantitative Gene Expression Fold model and the categorical Gene Expression Direction model. **(C)** Posterior class probabilities illustrating model confidence for correct (green) versus incorrect (red) predictions across the three clinical origins (CRC, Crohn's, Healthy colon). The horizontal dashed line indicates the theoretical threshold for random chance (0.33). **(D)** Gene-wise association with clinical disease class evaluated via Chi-squared test (left; expressed as -log10 *p*-value) and Cramér’s V (right). The red dashed line denotes the a priori significance threshold ($p = 0.001$), with significant associations highlighted in blue.

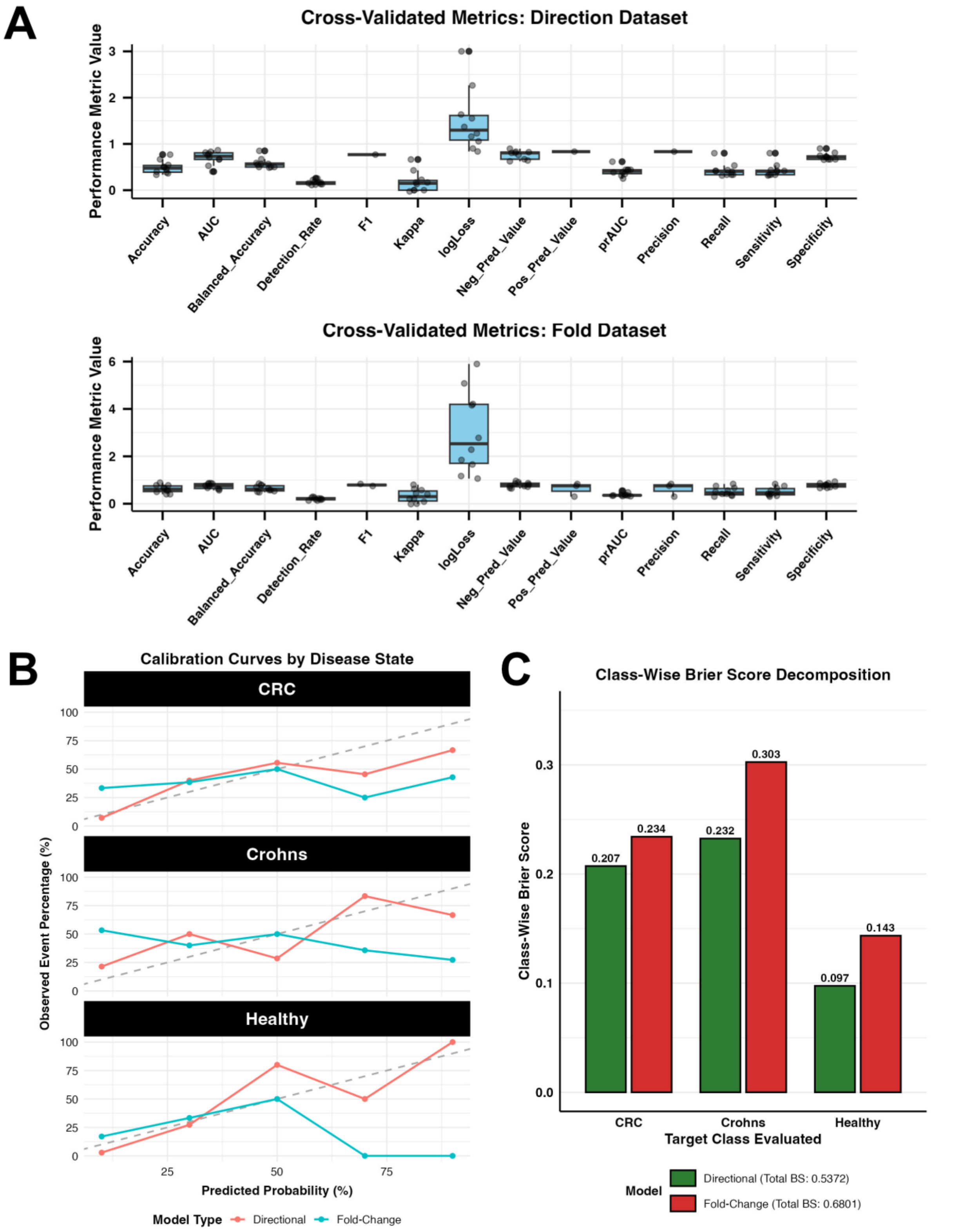


**Supplementary Figure 10: Cross-validation performance and calibration analysis of Naïve Bayes models. (A)** Boxplots summarizing performance metrics (Accuracy, AUC, F1, Kappa, logLoss, etc.) from 10-fold cross-validation for the Gene Expression

Direction (top) and Gene Expression Fold (bottom) models. Points represent individual fold results. **(B)** Calibration curves comparing predicted probability versus observed event percentage for the Directional (red) and Fold Change (blue) models across CRC, Crohn's, and Healthy classes. The dashed line indicates ideal calibration. **(C)** Class-wise Brier Score decomposition comparing the probabilistic accuracy of the Directional (green) and Fold-Change (red) models for each clinical origin. Lower Brier scores indicate better model calibration. Total Brier Scores are provided in the legend.
